# Data-driven detection of subtype-specific and differentially expressed genes

**DOI:** 10.1101/517961

**Authors:** Lulu Chen, Yingzhou Lu, Guoqiang Yu, Robert Clarke, Jennifer E. Van Eyk, David M. Herrington, Yue Wang

## Abstract

Tissue or cell subtype-specific and differentially-expressed genes (SDEGs) are defined as being differentially expressed in a particular tissue or cell subtype among multiple subtypes. Detecting SDEGs plays a critical rolse in molecularly characterizing and identifying tissue or cell subtypes, and facilitating supervised deconvolution of complex tissues. Unfortunately, classic differential analysis assumes a convenient null hypothesis and associated test statistic that is subtype-non-specific and thus, resulting in a high false positive rate and/or lower detection power with respect to particular subtypes. Here we introduce One-Versus-Everyone Fold Change (OVE-FC) test for detecting SDEGs. To assess the statistical significance of such test, we also propose the scaled test statistic OVE-sFC together with a mixture null distribution model and a tailored permutation scheme. Validated with realistic synthetic data sets on both type 1 error and detection power, OVE-FC/sFC test applied to two benchmark gene expression data sets detects many known and de novo SDEGs. Subsequent supervised deconvolution results, obtained using the SDEGs detected by OVE-FC/sFC test, showed superior performance in deconvolution accuracy when compared with popular peer methods.

---

Molecular characterization (e.g. gene expression profile) of a complex biologic system often includes features that are ubiquitously expressed by all cell or tissue types in the system (e.g. housekeeping genes)^1^, and expressed features that are specific to one or more cell or tissue subtypes present in the system (marker genes or differentially-expressed genes)^2-4^. An important but frequently underappreciated issue is how a “cell or tissue subtype-specific expression pattern” is defined. Ideally, a specific expression pattern would be composed of individual features that are specifically and differentially expressed in the cognate cell or tissue subtype of interest in relation to every others – so-called subtype-specifc and differentially expressed genes (SDEGs)^5-8^.

SDEGs play a critical role in molecularly characterizing and identifying tissue or cell subtypes, and facilitating the supervised deconvolution of complex tissues^5,8,9^. However, detecting SDEGs using tissue or cell-specific molecular expression profiles remains a challenging task^10^. For example, the most frequently used methods rely on an extension of ANOVA that identifies genes differentially expressed across any of the relevant cell or tissue subtypes. In this case, the null hypothesis is that samples in all subtypes are drawn from the same population, resulting in the selection of genes that may not conform to the SDEG definition. One-Versus-Rest Fold Change (OVR-FC) is another popular method based on the ratio of the average expression in a particular subtype to that of the average expression in the rest samples^10-12^, and OVR t-test is occasionally used to assess the statistical significance of detected genes^13^. However, a gene with low average expression in the rest is not necessarily expressed at a low level in every subtype in the rest. Expectedly, simulation studies show that Marker Gene Finder in Microarray data (MGFM) outperforms OVR t-test^14^. Alternative strategies include One-Versus-One (OVO) t-test and Multiple Comparisons with the Best (MCB)^15^ that use additional pairwise significance testing or the confidence intervals of OVO statistics^2,16^.

To address the critical problem of the absence of a detection method explicitly matched to the definition of SDEGs, here we introduce One-Versus-Everyone Fold Change (OVE-FC) test to detect SDEGs among many subtypes. OVE-FC test has previously been proposed to detect SDEGs to improve multiclass classification, where the selection is based on whether the mean of one subtype is significantly higher or lower than the mean from each of the other subtypes^5,6^. To assess the statistical significance of such test, we also propose the scaled test statistic OVE-sFC together with a mixture null distribution model. Because the expression patterns of non-SDEGs can be highly complex, a tailored permutation scheme is used to estimate the corresponding distribution under the null hypothesis.

We first validate the performance of OVE-sFC test on extensive simulation data, in terms of type 1 error rate and False Discovery Rate (FDR) control. We then demonstrate the detection power of the OVE-FC/sFC test over a comprehensive set of scenarios, in terms of partial area under the receiver operating characteristic curve (pAUC), and in comparison with top peer methods. We present the utility of OVE-FC/sFC test by applying it to benchmark public data, and assessing the performance by comparing with known SDEGs and by the accuracy of supervised deconvolution that uses the expression patterns of *de novo* SDEGs detected by OVE-FC/sFC test.

## Results

### Validation of OVE-sFC test on type 1 error using simulation data sets

To test whether our OVE-sFC test can detect SDEGs at appropriate significance levels, we assessed the type 1 error using simulation studies under the null hypothesis (Methods). Accuracy of type 1 error is crucial for any hypothesis testing methods that detect SDEGs based on their p-values because if the type 1 error is either too conservative or too liberal, the p-value is inflated by either too many false positive or false negative estimates, loses its intended meaning, and becomes difficult to interpret.

The simulation data contain 10,000 genes whose baseline expression levels are sampled from the real benchmark microarray gene expression data with replicates of purified cell subtypes (GSE19380^8^). Using realistic simulation data sets with various parameter settings, we show that in all scenarios the empirical type 1 error produced by OVE-sFC test closely approximates the expected type 1 error (**Figure 1a, Figure S2, Figure 2a-b**). The p-values associated with OVE-sFC test statistics exhibit a uniform distribution as expected. Moreover, even with unbalanced sample sizes among the subtypes, the mixture null distribution estimated by our posterior-weighted permutation scheme produces the expected empirical type 1 error rate (**Figure S2** and **Figure 2a**). In contrast, the empirical type 1 error produced by OVR t-test and OVO t-test either over-estimates or under-estimates the expected type 1 error. The p-values associated with OVR t-test and OVO t-test deviate from a uniform distribution (**Figure 1b**). We also evaluate the type 1 error associated with individual subtypes under high noise levels and small sample sizes. For each of the subtypes, experimental results show that the empirical type 1 error produced by OVE-sFC test closely matches the expected type 1 error (**Figure 1b** and Supplementary Information).

**Figure 1.**
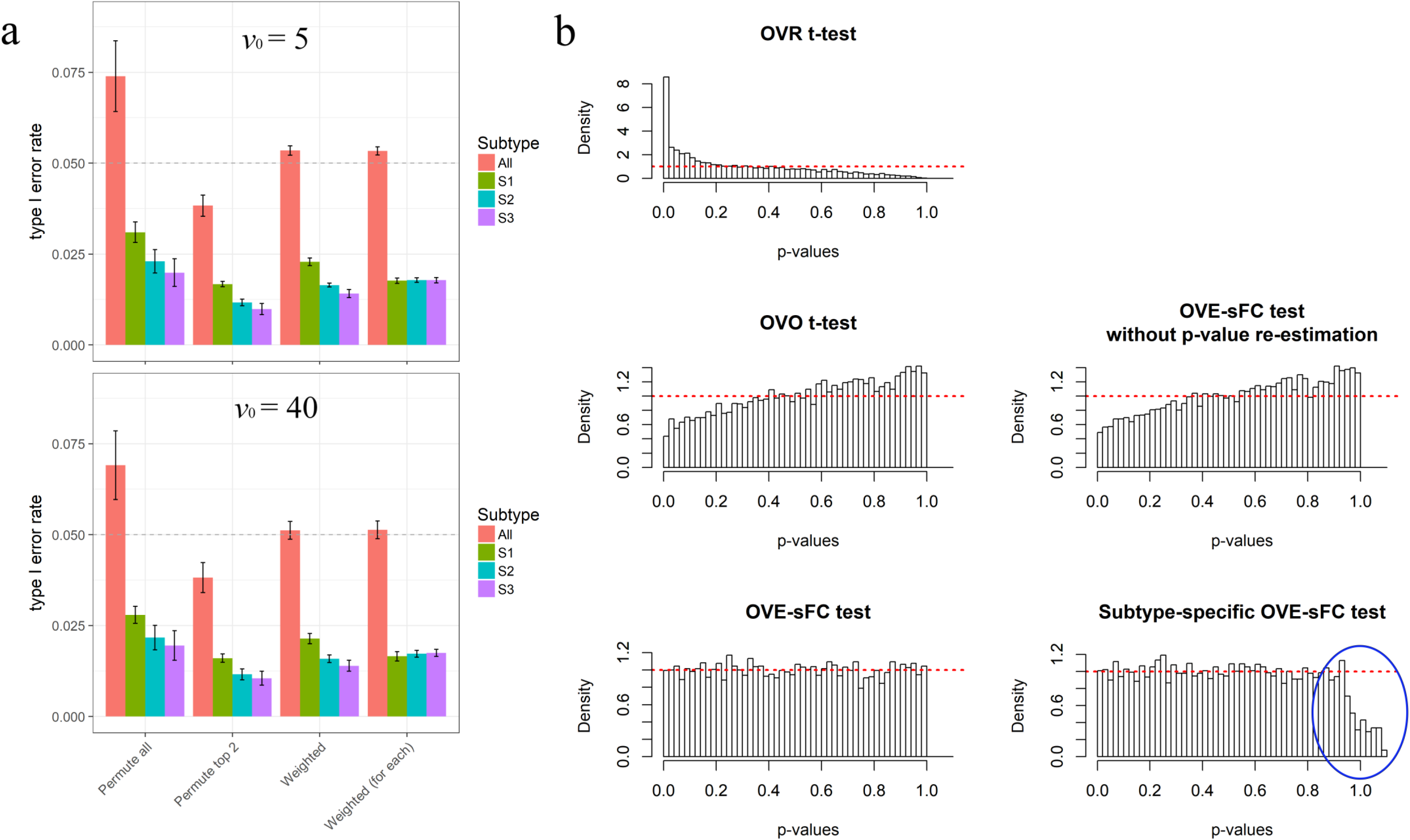
Assessment on Type 1 error rates and p-value distributions using simulated data sets under the null hypothesis, involving three subtypes with unbalanced sample sizes (*N*_1_ = 3, *N*_2_ = 6, *N*_3_ = 9). (a) Bar chart for the mean and 95% confidence interval of type I error rates with p-value cutoff at 0.05 over 30 simulated experiments, showing both overall and subtype-specific false-positive rates corresponding to different permutation schemes. (b) Histograms of p-value distributions associated with five SDEG detection methods, where simulation data consisted of 60% housekeeping genes, *σ*_0_ = 0.5 and *v*_0_ = 40. Note that subtype-specific p-values can be higher than 1.0 after multiple testing correction and thus will be truncated (indicated by the blue circle; see Supplementary Information for details).

**Figure 2.**
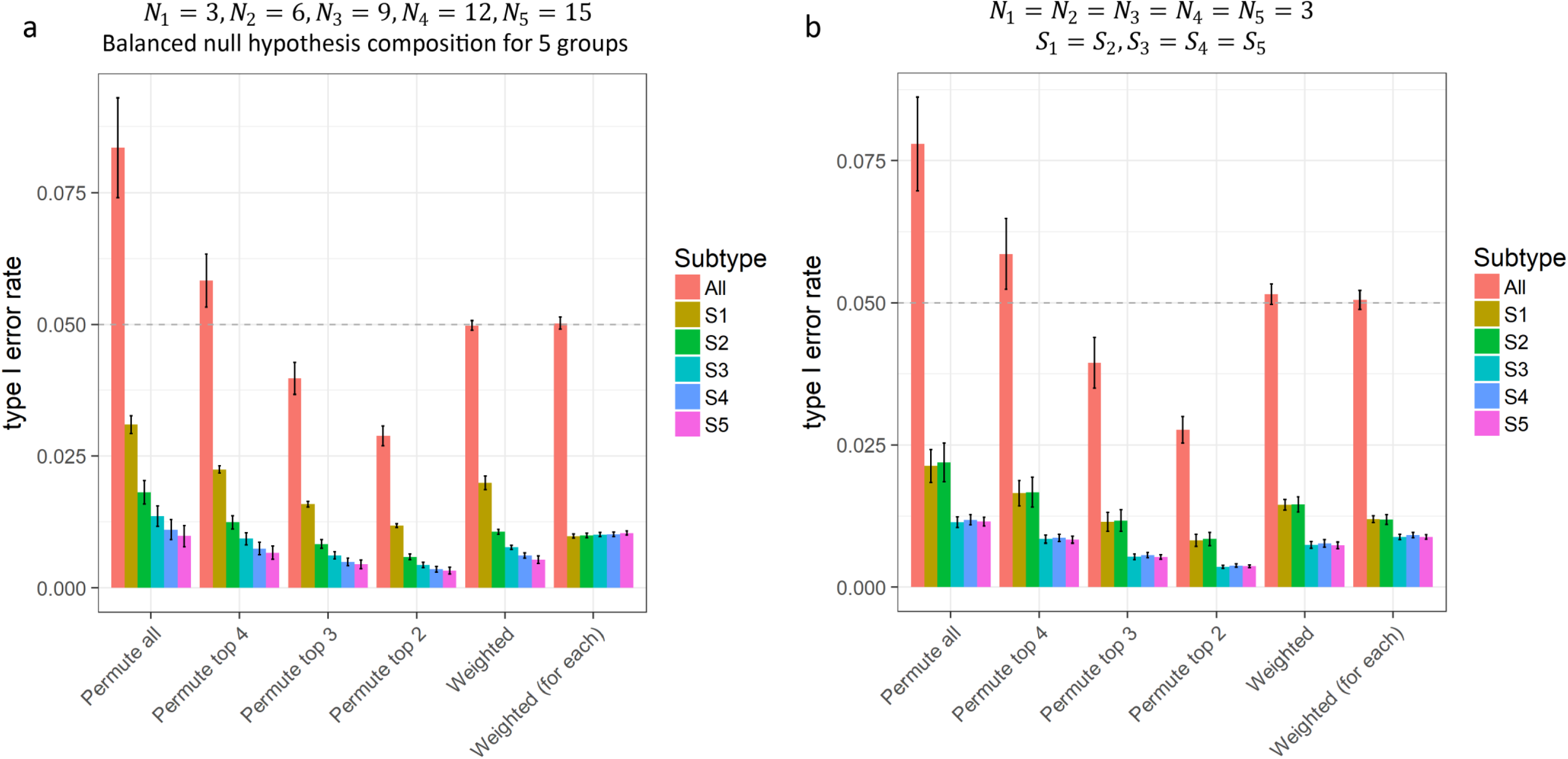
Assessment on Type 1 error rates using simulated data sets involving five subtypes. The results are obtained using the p-value cutoff at 0.05 over 30 experiments. (a) Bar chart of the mean and 95% confidence interval of type I error rates with unbalanced sample sizes. (b) Bar chart of the mean and 95% confidence interval of type I error rates with unbalanced compositions of mixture null distribution.

We conducted similar validation studies involving five subtypes over a wide range of simulation scenarios (**Error! Reference source not found.**). Experimental results again show that OVE-sFC test produces the empirical type 1 error rates that match the expected type 1 error rates. Furthermore, subtype-specific p-value estimates effectively balance the uneven type 1 error rates among the subtypes with different numbers of upregulated genes (Methods, **Figure 2b**, and Supplementary Information).

### Comparative assessment of OVE-FC/sFC test on power of detecting SDEGs using simulation data sets

Using realistic simulation data sets, we simulated a comprehensive set of scenarios to compare the power of OVE-sFC test and peer methods in detecting SDEGs (Method and Supplementary Information). Simulation data are generated again by modifying the expression levels of real gene expression data, where a portion of the genes are designated as SDEGs that are upregulated specifically in one of the participating subtypes, with fold change drawn in certain ranges (Supplementary Information). To recapitulate the characteristics of real expression data, various parameter settings are considered including unbalanced sample sizes or diverse mixture null distribution cross subtypes, each with 20 replications.

In large-scale multiple testing, False Discovery Rate (FDR) control is an important issue when assessing the detection power. For a well-designed significance test, the objective is to maximize power while assuring FDR below an allowable level. To test whether the q-value reflects the actual FDR, ‘fdrtool’ package is used to estimate the q-value for each gene^17^, where the empirical FDR with an estimated q-value of 0.05 is expected to be around 0.05. Another informative criterion is the pAUC that emphasizes the leftmost partial area under the receiver operating characteristic curve, focusing on the sensitivity at lower False Positive Rates (FPR)^18^.

Experimental results show that both overall and subtype-specific OVE-sFC test achieve a well-controlled FDR that matches the q-value cutoff (**Figure S5-S6**). In contrast, OVR t-test underestimates, while OVO t-test overestimates, the FDR (Supplementary Information). Moreover, subtype-specific OVE-sFC test exhibits a more balanced FPR for SDEGs across subtypes, while peer methods produce higher FPRs in the subtypes of small sample size.

For pAUC, the OVE strategy in OVE-FC/sFC test achieved the highest power in detecting SDEGs (**Figure 3, Figure S7-S8, Table S1-S3**). More specifically, for more realistic SDEGs (with sufficiently large fold change), OVE-sFC test shows the best performance; for the ideal SDEGs (i.e., marker genes with significantly large fold change^8,19^), both OVE-FC and OVE-sFC achieve the best performance with slight outperformance by OVE-FC. In comparison with the peer methods, OVE-sFC test consistently outperforms OVO t-test in these challenging experiments involving more subtypes and using RNAseq data. More importantly, the outperformance of OVE-FC/sFC over the peer methods at a stringent FPR range in ROC analysis is significant, because the related FDR is problematic in many real-world applications where large scale multiple comparisons are involved. In contrast, all three OVR methods exhibit lower detection power, and ANOVA has the lowest detection power (Supplementary Information).

**Figure 3.**
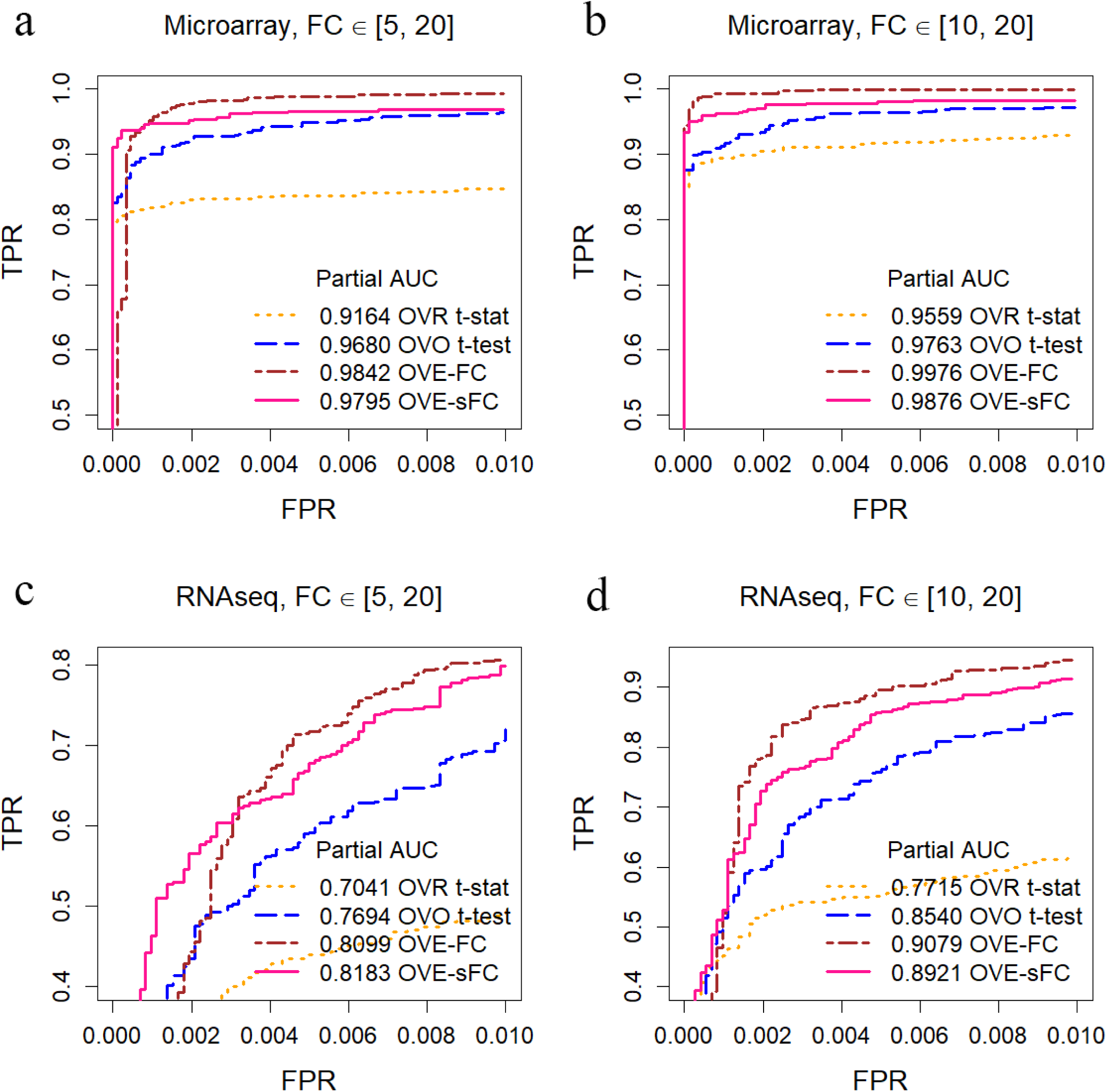
Assessment on detection power (partial ROC curves, FPR < 0.01) using real data derived simulations (data distribution is consistent with the base real dataset under null hypothesis; variances are sampled from real microarray data GSE28490 or RNAseq data GSE60424 with keeping mean-variance trend) involving seven unbalanced subtypes with various parameter settings. (a)(b) partial ROC curves across different FPR points on microarray-derived data. (c)(d) partial ROC curves across different FPR points on RNAseq-derived data. (OVR-FC and OVR t-test are not shown here due to low pAUC; subtype-specific OVE-sFC test’s performance is quite similar to OVE-sFC test; more complete ROC curves can be found in **Figure S7**.)

It is worth mentioning that when sample size is small, OVE-sFC test statistic borrows information across genes in estimating *a priori* variance via the limma method, thus stabilizing variance estimate for each gene. Furthermore, OVE-sFC test statistic estimates the parameters of the limma model from all subtypes, producing better results than that applying t-test independently with the limma model for each subtype pair. Indeed, for small sample size cases, our experimental results show that OVE-sFC test clearly outperforms OVO t-test (**Figure 3, Figure 5c** and **Tables S2-S3**). Note that when a large number of genes is involved, a more stringent multiple comparison correction or FPR/FDR control is applied.

**Figure 4.**
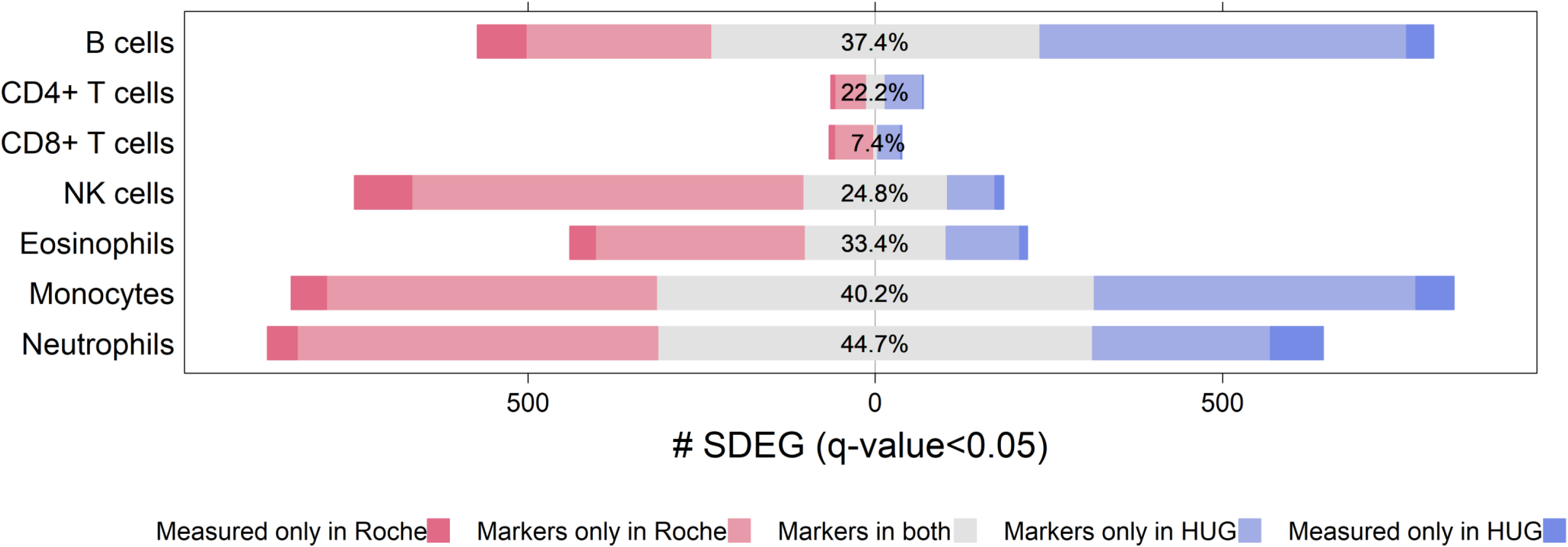
Percentile overlap of cell-type specific SDEGs between Roche and HUG datasets, quantified by Jaccard index (intersection over union). SDEGs are detected by subtype-specific OVE-sFC test with q-value < 0.05.

**Figure 5.**
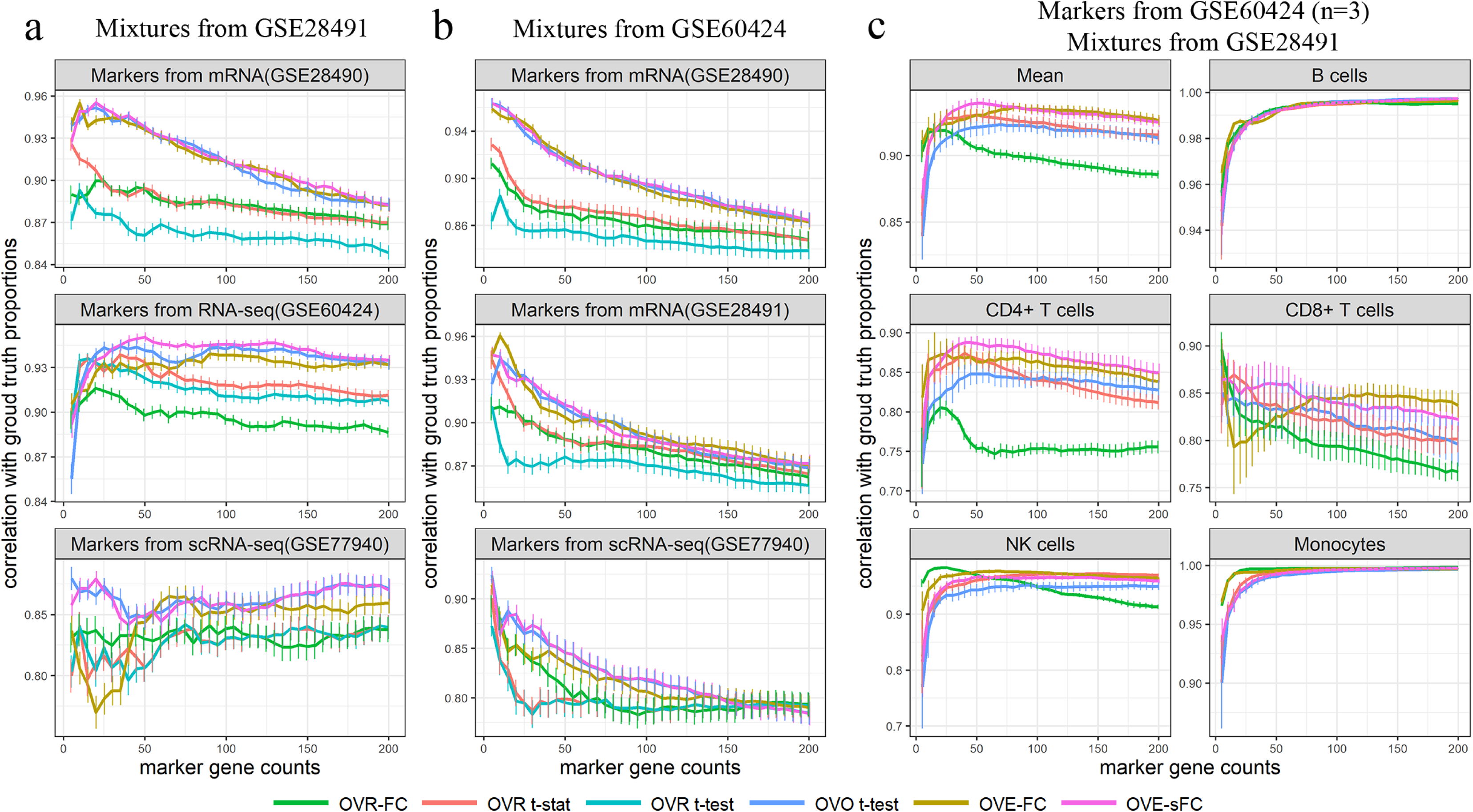
Correlation coefficients between CAM scores and ground truth proportions in simulated heterogeneous samples of mixed subtype mRNA expression profiles or RNAseq counts (a-c based on three different real gene expression datasets). CAM scores are estimated using the detected SDEGs from independent dataset and reflect the proportions of subtypes. The mean and 95% confidence interval are computed over 20 repeated experiments. (OVR t-test results are not shown in (c) due to very poor performance.)

### Application of OVE-sFC test on two benchmark gene expression data sets detects SDEGs (human immune cells)

We applied OVE-sFC test to two real microarray gene expression data sets, GSE28490 (Roche) and GSE28491 (HUG), to detect SDEGs associated with human immune cells^20^. In these data sets, the constituent subtypes are composed of seven different human immune cells that were isolated from healthy human blood: B cells, CD4+ T cells, CD8+ T cells, NK cells, monocytes, neutrophils, and eosinophils. Because Roche and HUG used the same protocols for cell isolation and sample processing from two independent panels of donors, the derived gene expression profiles enable use of a cross-validation strategy.

With an FDR control of q-value < 0.05 applied to both data sets, OVE-sFC test detects n=28 CD4+ T cell marker genes, n=7 CD8+ T cell marker genes, and multiple marker genes for other more distinctive cell types (**Table S4-S6**). Between the two data sets, we obtain a Jaccard index (intersection over union) of 36.8% for all SDEGs across all seven cell types. Overlap of monocyte and neutrophil marker genes detected from the two datasets is >40% (**Figure 4**). The number of SDEGs accounts for about one-third of all probesets (Roche: 39%, HUG: 34%). This result is expected because these subtypes are pure cell types and so more distinctive than would be seen with samples from multicellular complex tissues^9,21,22^. We also applied a Bonferroni multiple testing correction and a more stringent p-value < 0.001; the number of SDEGs account for 10.7% and 2.7% of all probesets in Roche and HUG data sets, respectively (**Table S4**), with only one common CD4+ T cell marker gene (FHIT) and one common CD8+ T cell marker gene (CD8B).

To present all kinds of combinatorial upregulation patterns among cell types occurred under the null hypothesis (**Figure S9**), probeset-wise posterior probabilities of component hypotheses in the null mixtures (Eq. 4) are accumulated and normalized to estimate the counterpart probabilities of the alternative hypotheses (Eq. S10), where the patterns of upregulation in B cells, monocytes, or neutrophils rank the top in both data sets, followed by upregulation in lymphoid cells (B cells, CD4+ T cells, CD8+ T cells, NK cells) and T cells (CD4+ T cells, CD8+ T cells) in the Roche dataset.

### Evaluation of ideal SDEGs detected by OVE-FC/sFC test via supervised deconvolution

Accurate and reliable detection of ideal SDEGs has significant impact on the performance of many supervised deconvolution methods that use the expression patterns of ideal SDEGs to score constituent subtypes in heterogeneous samples^21,23,24^. We adopted a Convex Analysis of Mixtures (CAM) score calculated from ideal SDEGs-guided supervised deconvolution to quantify the proportional aboundance of each subtype (Supplementary Information). The correlation coefficient between estimated scores and true proportions was used to assess the accuracy of various SDEGs selection methods.

Both OVE-FC and OVE-sFC tests are applied to three independent data sets acquired from the purified subtype expression profiles (GSE28490 Roche), purified subtype RNAseq profiles (GSE60424), and classified single-cell RNAseq profiles (GSE72056), respectively. The idea SDEGs are detected by six different methods including OVE-FC, OVE-sFC, OVR-FC, OVR t-stat, OVR t-test, and OVO t-test, and then used to supervise the deconvolution of realistically synthesized mixtures with ground truth.

The proportions of constituent subtypes are estimated by the CAM scores derived from expression levels of top-ranked SDEGs for each subtype. Supervised deconvolution results show that OVE-sFC test, OVE-FC test and OVO t-test methods achieved the highest correlation coefficients between CAM score and true proportions when compared with the performance of other methods (**Figure 5a, Figure S10**).

As a more biologically realistic case involving higher between-sample variations, we synthesize a set of n=50 *in silico* mixtures by combining the subtype expression profiles from bootstrapped samples in the RNAseq data set according to pre-determined proportions. Again, supervised deconvolution results show that the ideal SDEGs detected by OVE-FC test or OVE-sFC test or OVO t-test achieved superior deconvolution performance (**Figure 5b, Figure S11**).

As a more challenging case of RNAseq data with lower signal-to-noise ratio and small sample size, we repeated the simulations where *in silico* mixtures were synthesized by combining subtype mean expressions (GSE28491 HUG) and ideal SDEGs were detected from downsampled RNAseq profiles (GSE60424, n=3). Three purified samples were randomly selected for each subtype and analyzed by the six methods. The experimental results show that, in terms of ideal SDEG-guided deconvolution performance, OVE-sFC test strongly defeats OVO t-test. Moreover, OVE-sFC test outperforms OVE-FC test for phenotypically closer cell types (CD4+ T and CD8+ T cell types) (**Figure 5c).**

Across the varying number of ideal SDEGs (5∼200) being selected, **Figure 5** shows the impact of SDEGs (both at a fixed number and the corresponding content) selected by different methods on the performance of supervised deconvolution. While different subtypes are expected to have different number of ideal SDEGs practically and biologically, e.g., B cell or monocyte versus CD4+ T cell or CD8+ T cell, the fundamental working principle of various tissue deconvolution methods is that there is a proper small number of ideal SDEGs to be expressed exclusively in only one particular subtype. Thus applying a stringent OVE-sFC test p-value threshold, e.g., < 0.001 after correction (**Table S4**) is a good option, because suitable number of ideal SDEGs for CD4+ or CD8+ T cell is 5∼20, while B cells or monocytes often allows more ideal SDEGs to be used in supervised deconvolution.

## Discussions

Interpreting an expression profile of complex tissues requires both the knowledge of the relative abundance of the different cell or tissue subtypes and their individual expression patterns. Understanding the relative contribution of individual cell or tissue subtypes in individual samples may illuminate pathophysiologic mechanisms, biologic responses to various stimuli, or transitions in tissue phenotype - especially when the cell-cell and cell-matrix interactions in a complex system are necessary conditions for biological relevance. Expression patterns of SDEGs for the relevant cells or tissues can be used to support supervised deconvolution to estimate the relative prevalence of the contributing cell or tissue subtypes. Our present work is focused on SDEGs, *i.e.* restricted to the SDEG definition widely adopted^7,8,19,25^. Indeed, the work here is motivated by the need to obtain SDEGs for supervised *in silico* tissue deconvolution^21^ and/or tissue subtype characterization^9^, where the measured data are the mixtures of the expressions from multiple underlying subtypes and the SDEGs are used to estimate both the proportions of each subtype in individual heterogeneous samples as well as the averaged subtype-specific expression profiles.

Though ideal SDEGs are defined as being exclusively and consistently expressed in a particular tissue or cell subtype across varying conditions, biological reality dictates a more relaxed definition that allows SDEGs of a particular tissue or cell subtype having low or insignificant expressions in all other subtypes. Experimental results show that SDEGs detected by OVE-FC/sFC test with high thresholds or small p-values can accurately estimate both subtype proportions and expression profiles, serving as effective molecular markers (**Figure 5c, Figure S10 and S11**). Accuracy of OVE-FC/sFC test based SDEG detection may be affected by batch effect, normalization, and outliers; and reliability of OVE-sFC would depend on the variance estimate particularly when sample size is small.

OVE-sFC test makes a few assumptions and works best when all assumptions are valid. For example, while the proposed permutation scheme does not require the data to be normally distributed, under the null hypothesis, OVE-sFC test assumes that samples are drawn from the distribution of the same ‘shape’ for different genes, ensuring that the null distributions across genes can be aggregated together with variance-based standardization. Practically, when data distributions deviate significantly from a common shape, the limma-voom/vooma/voomaByGroup’s variance models can be used to accommodate unequal variances by appropriate observational-level weights^26^; when data distributions deviate significantly from normality, a permutational ANOVA can be used to estimate the null hypothesis components of the mixture distribution. Our experimental results show that with the mean-variance relationship estimated by limma-voom on RNASeq data, the OVE-sFC test can maintain the expected type 1 error rates or specified FDR (**Figure S6**). For outliers and drop-out zero values in RNAseq data, when needed, state-of-the-art two-group test methods designed specifically for RNAseq will be exploited and adopted, *e.g.* edgeR^27^, DESeq2^28^.

While OVE-FC is the simpler version of our OVE strategy and drives the newer OVE-sFC test, here we have demonstrated that OVE-FC is also an effective and robust method for detecting SDEGs particularly when sample size is small. On the other hand, OVE-sFC test is a critical complement to OVE-FC in twofold: (1) OVE-FC does not assess statistical significance (producing p-values) while OVE-sFC test specifically aims to provide a significance assessment and to potentially improve FDR control; (2) OVE-sFC test improves detection power in some of more challenging situations. Theoretically, detecting SDEGs by evaluating the significance with accurate p-values is an attractive feature of OVE-sFC test that can help control FDR at the expected level. Indeed, our experimental results show that the OVE-sFC test outperforms OVE-FC in the more challenging cases involving nonideal SDEGs (**Figure S8**) or phenotypically closer cell types (**Figure 5c**). Nevertheless, OVE-sFC test may become unstable when the scaling factor is too small or inaccurately estimated, and OVE-FC will not work well when pre-exclusion of extremely lowly-expressed genes is not done properly.

Though ANOVA has been the most commonly used method to test differences among the means of multiple subtypes, often in conjunction with a post-hoc Tukey HSD comparing all possible pairs of means^29^, it is not suitable for detecting SDEGs because the null hypothesis used by ANOVA does not truly enforce the definition of SDEGs. ANOVA detects differentially expressed genes rather than SDEGs and therefore produces too many false positives with respect to individual subtypes.

In addition to the SDEGs discussed here (genes uniquely up-regulated in a particular subtype), the counterpart subtype-specific down-regulated genes (genes uniquely down-regulated in a particular subtype) are also of biological interest^5^. OVE-FC/sFC test is principally applicable to detecting down-regulated SDEGs by reversing the comparison rule ^5^. There are certainly other alternative definitions of ‘informative genes’ for different analytical purposes, *e.g.*, sample classification. In our earlier work on multiclass classification^5,6^, we have shown that upregulated SDEGs selected by OVE-FC are sufficient to achieve multiclass classification and can often improve classifier performance over alternative informative gene subsets of the same size.

Lastly, when subtype-specific expression patterns are unknown, unsupervised deconvolution techniques (e.g., CAM^21^) are required. A theoretical advantage of unsupervised deconvolution is that it can identify both the cell/tissue subtype proportions and their specific expression patterns, albeit with possibly less fidelity than when neither is known a priori or measured from the same sample.

## Methods

### Basic OVE-FC test

Consider the measured expression level *s*_*k*_(*i, j*) of gene *j* in sample *i* across *k* = 1, …, … *K* subtypes. We denote the mean and variance of the logarithmic expression levels log *s*_*k*_(*i, j*) of gene *j* in subtype *k* by *μ*_*k*_(*j*) and *σ*^2^(*j*), respectively. Accordingly, OVE-FC test after logarithm for gene *j* is defined as the gap between the highestly expressed subtype and the second highestly expressed subtype ^5,14^,

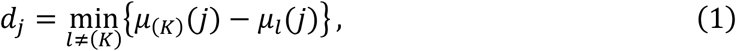

where subscript (*K*) indicates the subtype with the maximum mean among all subtypes. Note that OVE-FC has previously been proposed for multiclass classification^13, 14^, and matches well the definition of SDEGs ^5,8,19,25^. Conceptually, the null hypothesis for non-SDEGs, and the alternative hypothesis for SDEGs, can be described as

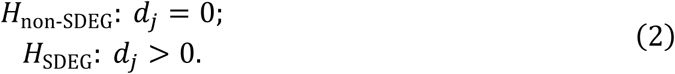

It is worth reiterating that the ideal SDEGs detected by OVE strategy with a stringent threshold are also termed the marker genes for supervised deconvolution^8,19^, and are principally similar to what detected by the Convex Analysis of Mixtures (CAM) method for fully unsupervised deconvolution^9,21^, i.e. the marker genes resided near the vertices of the scatter simplex.

### OVE-sFC test statistic and null distribution modeling

To assess statistical significance of OVE-FC and to cross-fertilize the information among genes, we assume that log *s*_*k*_(*i, j*) ∼*N*(*μ*_*k*_(*j*), *σ*^2^(*j*)), and further define the scaled test statistic OVE-sFC as

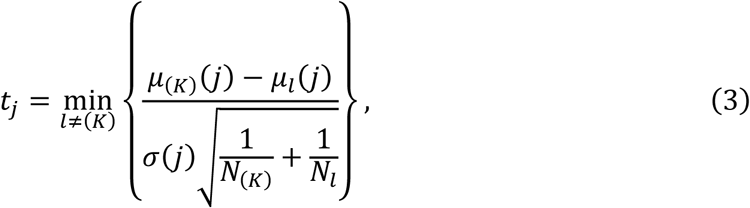

where *N*_(*K*)_ and *N*_*l*_ are the numbers of samples in subtypes (*K*) and *l*, respectively. However, for more than two subtypes *K* ≥ 3, modeling the distribution of *t*_*j*_ under the null hypothesis is challenging because the expression patterns of non-SDEGs are highly complex, i.e., non-SDEGs include both housekeeping genes and various combinatorial forms of differentially-expressed genes among the subtypes.

We propose the following mixture distribution of OVE-sFC test statistic *t* under the null hypothesis (**Figure 6**)

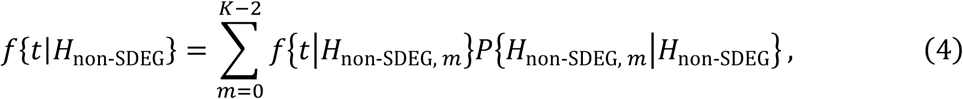

where *H*_non-SDEG, *m*_ is the *m*th component of the mixture null hypothesis *H*_non-SDEG_. We design a novel nested permutation scheme that approximates the complex null distribution and is consistent with the definition of SDEGs. Principally, *H*_non-SDEG, *m*_ is constructed by permuting the samples in the top (*K* − *m*) subtypes with higher mean expressions; that is, the samples in the bottom *m* subtypes with lower mean expressions are removed from the permutation. Note that *H*_non-SDEG, 0_ corresponds to the same null distribution used in ANOVA where all samples participate in the permutation.

**Figure 6.**
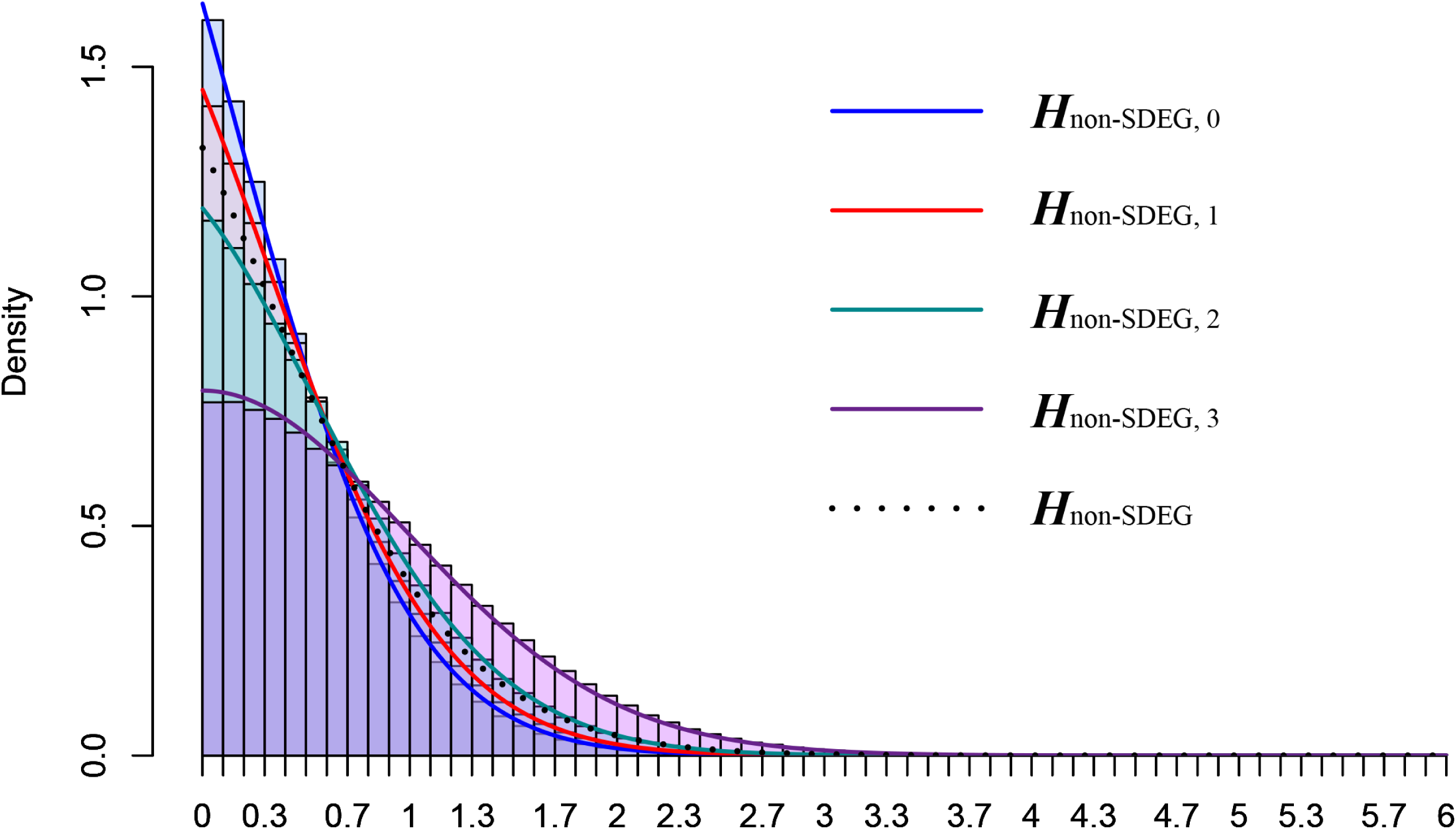
Mixture null distribution of OVE-sFC test statistics for detecting SDEGs. The mixture distribution consists of (*K* − 1) null components, each estimated from permuting samples in the top (*K* − *m*) subtypes of high mean expressions and weighted by the posterior probabilities of component null hypotheses.

This mixture null distribution model is proposed to model unknown yet potentially complex expression patterns of non-SDEGs under the null hypothesis. Accordingly, the specifically-designed permutation scheme(s) estimates such a mixture null distribution. The main advantage of the proposed permutation scheme(s) is its flexibility and comprehensiveness, which well match the mixture null distribution of various types and combinations. With varying the proportions of different non-SDEG types, OVE-sFC test is able to maintain the type 1 error rate close to the expected level with the help of the proposed permutation scheme(s) and the conditional probability of each non-SDEG type (**Figure S2**, Supplementary Information).

Note that *H*_non-SDEG,*m*_, *m* = 0, …, *K* − 2 represents (*K* − 1) different null hypotheses, each with an individualized null distribution that can be estimated by specific permutation scheme(s), *e.g.*, permuting samples in the top (*K* − *m*) subtypes. Collectively, a mixture of null distribution is constructed via combinations of different null hypotheses in various proportions. In contrast, without conditioning on *H*_non-SDEG,*m*_, all null distributions are aggregated equally into the mixture null distribution in the same proportion. Consequently, this simpler permutation scheme produces an equal-weight mixture model that cannot represent the complexity of the null distribution. Thus, the null distribution of the OVE-sFC test statistic could become distorted, *e.g.*, a uniform distribution of p-values in null data is not guaranteed, and the observed False Discovery Rate may not match the expected level.

Specifically, the null distribution of OVESEG-test statistics under *H*_non-SDEG, *m*_ is estimated from permuted samples and aggregated from different genes with weights. Let ***s***(*j*) = [*s*(1, *j*), …, *s*(*N, j*)] denote the measured expression vector of gene *j* across samples, where *N* is the total number of samples. These weights are the posterior probabilities of a component null hypothesis given the observation Pr{*H*_non-SDEG, *m*_|***s***(*j*)}, estimated by the local FDR fdr_non-SDEG, *m*_(*j*) ^30^, given by

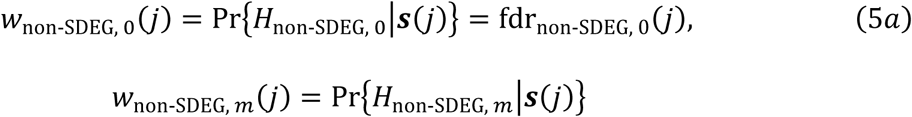

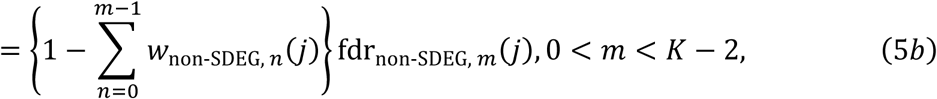

where fdr_non-SDEG, 0_(*j*) is the local FDR associated with ANOVA on all subtypes, and fdr_non-SDEG, *m*_(*j*) is the local FDR associated with ANOVA on the top (*K* − *m*) subtypes, estimated using R package “fdrtool” ^17^ (Supplementary Information).

### Assessing statistical significance of candidate SDEGs

The p-values of candidate SDEGs are estimated using the learned ‘mixture’ null distribution

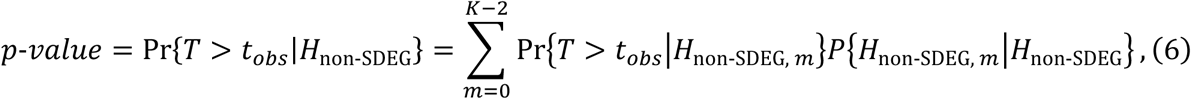

where *t*_*obs*_ is the observed OVE-sFC test statistic, and *T* is the continuous dummy random variable. Specifically, Pr{*T* > *t*_*obs*_|*H*_non-SDEG, *m*_} is calculated by the weighted permutation scores

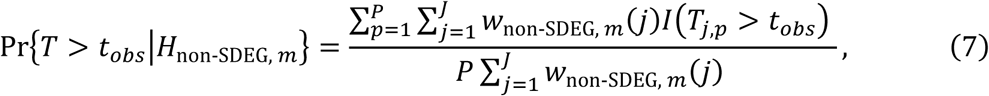

where *P* is the number of permutations, *J* is the number of participating genes, *I*(.) is the indicator function, and *T*_*j,p*_ is the OVE-sFC test statistic in the *p*th permutation on *j*th gene. Furthermore, the component weight in the mixture null distribution is estimated by the membership expectation of the posterior probabilities over all genes

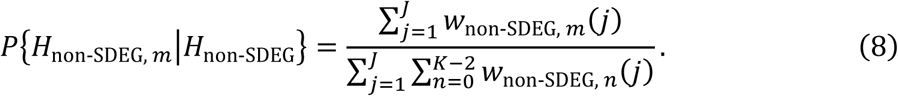

Lastly, substituting (7) and (8) into (6), the p-value associated with gene *j* is calculated by:

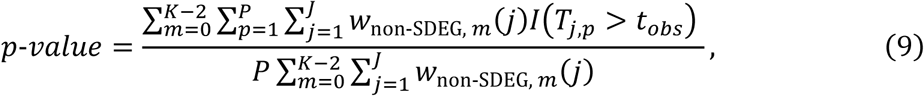

with a lower bound of min 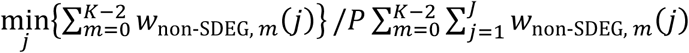.

Supplementary Information provides more details on the deviation of OVE-sFC test p-values when considering all subtypes together (Eq. 9) and when considering one subtype specifically (Eq. S7-S8).

### Empirical Bayes moderated variance estimator of within-subtype expressions

The importance of an accurate estimator on pooled within-subtype variance *σ*^2^(*j*) is twofold - calculating the OVE-sFC test statistic *t*_*j*_ and determining the local false discovery rate fdr_non-SDEG, *m*_(*j*), particularly with small sample size. We assume a scaled inverse chi-square prior distribution 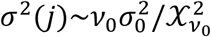, where *v* and 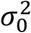 are the prior degrees of freedom and scaling parameter, respectively^31^. We then adopt the empirical Bayes moderated variance estimator 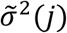 that leverages information across all genes, as used in *limma* and given by

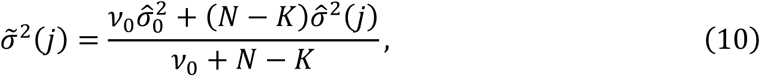

where *N* is the total number of samples, and 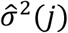 is the pooled variance estimator, given by

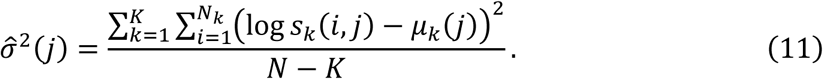

The prior parameters *v*_0_ and 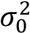 are estimated from the pooled variances. The moderated variances shrink the pooled variances towards the prior values depending on the prior degrees of freedom and the number of samples. Note that *t*-*stat*(*j*) with moderated variance estimator 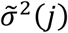 follows a *t*-distribution with *v*_0_ + *N* − *K* degrees of freedom (Supplementary Information).

### Brief review of the most relevant peer SDEG selection methods

The OVR-FC uses a simple test defined by

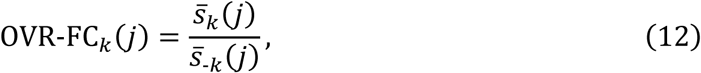

where 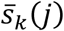 and 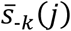 are the geometric means of the *j*th gene expressions within subtype *k* and associated with the combined remaining subtypes, respectively. The OVR t-test uses a statistical test given by

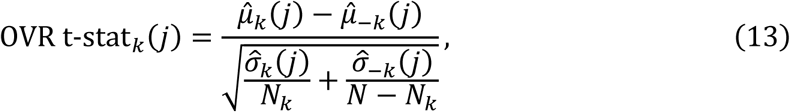

where 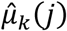 and 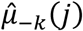 are the sample means of the *j*th gene expressions within subtype *k* and associated with the combined remaining subtypes, respectively; and 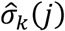 and 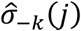 are the sample variance of the *j*th gene expressions within subtype *k* and associated with the combined remaining subtypes, respectively. The OVO t-test, conducts t-tests among all subtype pairs and selects genes upregulated in one subtype for all the involved tests, where the variances are estimated only from every pair of subtypes^16^ (Supplementary Information). In contrast, the OVE-sFC test exploits all subtypes in estimating the variances. The benefit of using all subtypes for modeling is significant in challenging cases with higher variance, smaller sample size, and more subtypes (Supplementary Information).

### Simulation study for validating OVE-sFC test statistics on type 1 error

Among the 10,000 simulated genes, a portion are housekeeping genes that take the baseline expression levels across all subtypes under *H*_non-SDEG, 0_. Expression levels of the remaining genes are adjusted to exhibiting similar upregulations in at least two subtypes, mimicking all types of participating null hypotheses. The upregulations are modeled by uniform distribution(s) in scatter space, with variance following an inverse chi-square distribution 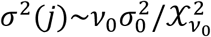, where the prior degree of freedom *v*_0_ takes 5 or 40, and *σ*_0_ takes 0.2, 0.5, or 0.8 (Supplementary Information).

### Gene expression data of human immune cells (GSE28490 and GSE28491)

In these data sets, each cell subtype consists of at least five samples, excluding few outliers (**Table S7**). Following preprocessing of the raw measurements, 12,022 probesets in Roche and 11,339 probesets in HUG were retained and used in the analyses (Supplementary Information).

### Realistic synthetic data for supervised deconvolution

Five subtypes (B cell, CD4+ T cell, CD8+ T cell, NK cell, monocytes) were included in synthesizing n=50 *in silico* mixtures, where purified subtype mean expressions (GSE28491 HUG) were combined according to pre-determined proportions with additive noise, simulating heterogeneous biological samples (Supplementary Information).

### Availability of package and supporting data

A Bioconductor approved R package of OVE-sFC test is freely available at http://bioconductor.org/packages/OVESEG. A detailed user’s manual and a vignette are provided within the package. In addition, public gene expression data analyzed in this paper are also available from the Gene Expression Omnibus Database under Accession Number GEO: GSE19380, GSE28490, GSE28491, GSE60424, and GSE72056.

## ADDITIONAL INFORMATION

Supplementary information accompanies this paper at https://doi.org/

## ACKNOWLEDGMENTS

This work was funded in part by the National Institutes of Health under Grants HL111362-05A1, HL133932, W81XWH-18-1-0723 (BC171885P1), and U01NS115658-01.

## AUTHOR CONTRIBUTIONS

L.C. and Y.W. developed OVE-FC/sFC test framework and wrote the manuscript; L.C. implemented OVE-FCs/FC test algorithm; L.C and Y.L. performed real data analysis; D.M.H., J.E.V.E. and R.C. interpreted results and edited the manuscript; G.Y. provided statistical expertise support.

## COMPETING FINANCIAL INTERESTS

The authors declare no competing financial interests.

