## Supplementary Information for "Data-driven detection of subtype-specific and differentially expressed genes"

[1]Department of Electrical and Computer Engineering, Virginia Polytechnic Institute and State University, Arlington, VA 22203, USA; [2]Lombardi Comprehensive Cancer Center, Georgetown University, Washington, DC 20057, USA; [3]Advanced Clinical Biosystems Research Institute, Cedars Sinai Medical Center, Los Angeles, CA 90048, USA; [4]Department of Internal Medicine, Wake Forest University, Winston-Salem, NC 27157, USA

[*]Author for correspondence: Yue Wang, Ph.D.
Virginia Tech Research Center - Arlington
900 N. Glebe Road, Arlington, VA 22203


### OVE-sFC test

#### OVE-sFC test statistic

Motivated by our earlier work on OVE-FC[1], we define One-Versus-Everyone Log Fold Change (OVE-LFC) as

$$d_{jk} = \mu_k(j) - \max_{l \neq k} \mu_l(j) = \min_{l \neq k}\{\mu_k(j) - \mu_l(j)\}, k = 1, \dots, K, \qquad (S1)$$

where $\mu_k(j)$ is the mean of logarithmic expressions of gene $j$ in subtype $k$. Conceptually, the the null hypothesis for non-SDEGs of subtype $k$ and alternative hypothesis for SDEGs of subtype $k$ can be described as

$$\begin{aligned} H_{non\text{-}MG(k)}&: d_{jk} \leq 0; \\ H_{MG(k)}&: d_{jk} > 0; \end{aligned} \qquad (S2)$$

Standardizing OVE-LFC by variance, we obtain the subtype-specific OVE-sFC test statistic

$$t_{jk} = \min_{l \neq k} \left\{ \frac{\mu_k(j) - \mu_l(j)}{\sigma(j)\sqrt{\frac{1}{N_k} + \frac{1}{N_l}}} \right\} = \min_{l \neq k}\{t\text{-}stat_{k,l}(j)\}, k = 1, \dots, K, \qquad (S3)$$

where $\sigma^2(j)$ is genewise variance of logarithmic expressions within one subtype, $t\text{-}stat_{k,l}(j)$ is t-statistic between subtype $k$ and $l$. Using $t_{jk}$ to select SDEGs for subtype $k$, denoted by $\text{SDEG}(k)$, is equivalent to performing pairwise t-tests between subtype $k$ and each of the remaining subtypes and then taking their intersections, since the the null hypothesis and alternative hypothesis can be rewritten as $H_{\text{non-SDEG}(k)} = \bigvee H_{\text{non-SDEG}(k,l)}$ and $H_{\text{SDEG}(k)} = \bigwedge H_{\text{SDEG}(k,l)}$ respectively, where $\text{SDEG}(k, l)$ denotes SDEGs enriched in subtype $k$ against subtype $l$. However, naively combining the pairwise t-test p-values to assess the significance of SDEGs did not model the the null distributions rigorously according to the definition of SDEGs.

Notice that we only need to compute $t_{jk}$ when $\mu_k(j)$ is the largest value among $\mu_l(j), l = 1, \dots, K$, as we only need to determine whether the highest expressed subtype has significant enrichment against other subtypes. Thus, we can define OVE-sFC test statistic, regardless of a specific subtype, as

$$t_j = \max_{k=1,\dots,K}\{t_{jk}\} = t_{j(K)} \quad (S4)$$

to test the significance level of gene $j$ as a SDEG (**Eq. 2 and Eq. 3**), where subscript $(K)$ indicates the $K$th order of ranked sequence $[\mu_l(j), l = 1, \dots, K]$, i.e. the subtype with maximum mean.

#### Difficulty to assess statistical significance of candidate SDEGs

To evaluate the significance level of SDEGs, we need to estimate the distribution of OV-sFC test statistics conditioned on the the null hypothesis $H_{\text{non-SDEG}}: d_j = 0$, which includes the following $(K-1)$ the null hypotheses (**Fig. S1a**):

$$H_{\text{non-SDEG, }0}: \mu_{(K)}(j) = \mu_{(K-1)}(j) = \cdots = \mu_{(1)}(j);$$

$$H_{\text{non-SDEG, }1}: \mu_{(K)}(j) = \mu_{(K-1)}(j) = \cdots = \mu_{(2)}(j) > \mu_{(1)}(j);$$

...

$$H_{\text{non-SDEG, }K-2}: \mu_{(K)}(j) = \mu_{(K-1)}(j) > \mu_{(K-2)}(j), \dots, \mu_{(1)}(j);$$

where subscript $(k)$ enclosed in parentheses indicates the $k$th order of $[\mu_l(j), l = 1, \dots, K]$. The the null distribution of OVE-sFC test statistics under $H_{\text{non-SDEG, }m}$, $0 < m \le K-2$, is asymptotically equivalent to that under $H'_{\text{non-SDEG, }m}: \mu_{(K)}(j) = \cdots = \mu_{(m+1)}(j) \gg \mu_{(m)}(j), \dots, \mu_{(1)}(j)$ when $\Delta_m(j) = \mu_{(K)}(j) - \mu_{(m)}(j)$ is sufficiently large and thus only the highest expressed $(K-m)$ subtypes affect OVE-sFC test statistics. On the contrary, when $\Delta_m(j)$ approaches zero, $H_{\text{non-SDEG, }m}$ will tend to become $H_{\text{non-SDEG, }m-1}$ with the the null distribution of OVE-sFC test statistics becoming less dispersed, as shown in the following simulation.

To illustrate the change of the null distribution with varied $\Delta_m(j)$, we can calculate theoretical p-values based on a multivariate normal/student's t-distribution of $(K-1)$ random variables $t\text{-}stat_{(K),l}, l = 1, \dots, K, l \ne (K)$ which have mean vector $[\mathbf{0}_{K-m-1}, \Delta_m(j), \dots, \Delta_1(j)]$, unit variances, and correlation coefficients

$$\rho_{l_1,l_2} = \frac{1}{\sqrt{\left(\frac{N_{(K)}}{N_{l_1}}+1\right)\left(\frac{N_{(K)}}{N_{l_2}}+1\right)}}. \qquad (S5)$$

Decreasing $\Delta_m(j)$ tends to shift the distribution of $\min_{l \neq (K)}\{t\text{-}stat_{(K),l}\}$ in the negative direction, increasing the significance level (with smaller p-values). For example, when $K = 3$ with an equal sample size for all subtypes, the correlation coefficient between $Z_0 = t\text{-}stat_{(3),(2)}$ and $Z_1 = t\text{-}stat_{(3),(1)}$ is $\rho = 0.5$. Under the null hypothesis $H_{\text{non-SDEG},1}$, $Z_0$ has zero mean and $Z_1$ could have a positive mean. **Fig. S1b** shows the change of p-values with varying $Z_1$. If the mean of $Z_1$ is large, three-group OVE-sFC test statistics can be approximated by two-group t-statistics, yielding the same significance level as a t-test. Otherwise, the p-value will decrease and reach as low as 0.01387 when the mean of $Z_1$ also equals zero.

In fact, if we know which the null hypothesis each gene comes from as well as the true $\Delta_m(j)$, we can calculate the theoretical p-value. Of course, these are unknown and have to be modeled and approximated. We propose a mixture model of the null distributions under the component hypotheses (**Eq. 4-5)** to estimate the p-values of candidate SDEGs (**Eq. 6-9**).

#### Mixture model to assess statistical significance of candidate SDEGs

This subsection gives a detailed explanation of **Eq. 4-9**.

(1) The order of subtypes

When $H_{\text{non-SDEG},m}$ is true, samples in the highest expressed $(K - m)$ subtypes are assumed to be drawn from the same populations. Hence, we need to identify which $(K - m)$ subtypes are the highest expressed. To obtain meaningful OVE-sFC test statistics (**Eq. 3**), we set the highest expressed subtype to be the one with the largest mean and sort the remaining by their t-statistics against the highest expressed subtype. The order of subtypes is exactly equal to the order of $[\mu_l(j), l = 1, \dots, K]$ when $N_1 = \cdots = N_K$. The subtype with the largest estimated mean does not always have the largest true population mean, due to high variance, especially when sample sizes are unbalanced among all subtypes. Nonetheless, we can focus on the significance level of SDEGs for the subtype with the largest estimated mean. While a gene may be expressed highest in one of the remaining subtypes, and thus correspond to a more significant level in other subtypes, this gene is not likely to be an adequate marker with a low p-value.

(2) $\mathrm{f}\{t|H_{\text{non-SDEG}, m}\}$

We approximate $H_{\text{non-SDEG}, m}$ by $H'_{\text{non-SDEG}, m}$ under which the the null distribution is estimated by permuting sample labels in the top $(K-m)$ subtypes. These permutation distributions from different genes could be pooled together with certain weights where: **(a)** the gene with large posterior probabilities of $H_{\text{non-SDEG}, m}$ should contribute more to the the null distribution under $H_{\text{non-SDEG}, m}$ and **(b)** genes with relatively small $\Delta_{jm}$ may affect the the null distribution estimation under both $H_{\text{non-SDEG}, m}$ and $H_{\text{non-SDEG}, m-1}$. As local FDR is a good estimator of the posterior probability of a the null hypothesis [2], the weight of each gene contributing to a component the null hypothesis is assigned according to the local FDR of ANOVA on the observed expression values across certain subtypes (**Eq. 5**). Note that we include the genes in the $fdr_{\text{non-SDEG, m}}$ calculation with a probability of $\{1-\sum_{n=0}^{m-1} w_{\text{non-SDEG}, n}(j)\}$ so that the computation of $w_{\text{non-SDEG}, m}$ is less affected by genes associated with $H_{\text{non-SDEG}, 0}, \dots, H_{\text{non-SDEG}, m-1}$. R package "fdrtool" [3] is then used to estimate local FDR. The the null distribution of OVE-sFC test statistics under $H_{\text{non-SDEG}, m}$ are approximated by weighted permutations (**Eq. 7**).

(3) $P\{H_{\text{non-SDEG}, m}|H_{\text{non-SDEG}}\}$

Given the posterior probabilities of all genes (**Eq. 5**), the component weight in the mixture the null distribution are estimated directly by

$$\begin{aligned} P\{H_{\text{non-SDEG}, m}|H_{\text{non-SDEG}}\} &= \frac{\Pr\{H_{\text{non-SDEG}, m}\}}{\sum_{n=0}^{K-2} \Pr\{H_{\text{non-SDEG}, n}\}} \\ &= \frac{\frac{1}{J}\sum_{j=1}^{J} \Pr\{H_{\text{non-SDEG}, m}|\boldsymbol{s}(j)\}}{\frac{1}{J}\sum_{j=1}^{J}\sum_{n=0}^{K-2} \Pr\{H_{\text{non-SDEG}, n}|\boldsymbol{s}(j)\}} \\ &= \frac{\sum_{j=1}^{J} w_{\text{non-SDEG}, m}(j)}{\sum_{j=1}^{J}\sum_{n=0}^{K-2} w_{\text{non-SDEG}, n}(j)}, \end{aligned} \tag{S6}$$

i.e. **Eq. 8**.

(4) p-values of candidate SDEGs

The final equation for OVE-sFC test p-values, **Eq. 9**, depends on $PJ(K-1)$ permutations and $J(K-1)$ ANOVA tests. As the observations are also included as one random permutation, OVE-sFC test p-value has a lower bound of $\min_j\left\{\sum_{m=0}^{K-2} w_{\text{non-SDEG},\,m}(j)\right\}/P\sum_{m=0}^{K-2}\sum_{j=1}^{J} w_{\text{non-SDEG},\,m}(j)$.

#### Subtype-specific p-values of candidate SDEGs

Using $t_j = \max_{k=1,\ldots,K}\{t_{jk}\}$ as a test statistic in K-group comparison is equivalent to using the absolute value of the statistic in the two-group differential test, which gives a concise way to define extremes in the $K$ directions. However, in the two-group differential test, if the test statistics have an asymmetric the null distribution for two sides, another widely-accepted choice is to double the smaller of the two tail regions[4, 5]. Extending this method to $K > 2$ cases, we can calculate subtype-specific p-values based on the the null distribution of each $t_{jk}, k = 1, \ldots, K$, by

$$\begin{aligned} p\text{-}value(k) &= \Pr\{T_k > t_{obs,k} \mid H_{\text{non-SDEG},\,m}\} \\ &= \sum_{m=0}^{K-2} \Pr\{T_k > t_{obs,k} \mid H_{\text{non-SDEG},\,m}\} P\{H_{\text{non-SDEG},\,m} \mid H_{\text{non-SDEG}}\} \\ &= \frac{\sum_{m=0}^{K-2}\sum_{p=1}^{P}\sum_{j=1}^{J} w_{\text{non-SDEG},\,m}(j) I(T_{jk,p} \geq t_{k,obs})}{P\sum_{m=0}^{K-2}\sum_{j=1}^{J} w_{\text{non-SDEG},\,m}(j)}, \end{aligned} \qquad (S7)$$

as one-tailed p-values. We then multiply the smallest tail by K. $t_{k,obs}$ is the observed OVE-sFC test statistic for a gene being tested under subtype $k$, and $T_{jk,p}$ is OVE-sFC test statistic in the $p$th permutation on $j$th gene under subtype $k$. We also need to avoid the extra computational burden brought about by calculating OVE-sFC test statistic for every subtype. Thus, a gene's p-value is calculated only for the subtype with highest mean, $p\text{-}value((K))$, corresponding to a positive $t_{k,obs}$. The OVE-sFC test statistics of permutated samples, $T_{jk,p}$, are also calculated only for the subtype with the highest mean. The OVE-sFC test statistics for other subtypes must be negative and could be set to be any negative constant because $t_{k,obs} > T_{jk,p}$ always holds in this case. Then the multiple-tailed p-value becomes

$$p\text{-}value = \min_{k=1,\ldots,K}\{p\text{-}value(k)\} * K \doteq p\text{-}value((K)) * K. \qquad (S8)$$

The right-hand side of **Eq. S8** does not always hold, resulting in p-values > 1, which is also a drawback of tail-doubling for two-tailed p-values. However, p-values for subtypes other than $(K)$ are unimportant. Even if the smallest tail appears under other subtypes, this gene is less likely to be an adequate SDEG. "Multiplying a one-sided p-value by K" can also be explained as multiple testing correction. Each molecule is actually tested $K$ times but we only record the p-value associated with the highest expressed subtype.

Subtype-specific OVE-sFC test p-values differ from the overall p-values in **Eq. 9** when the OVE-sFC test statistic is asymmetric across subtypes due to unbalanced sample sizes and/or unbalanced the null hypothesis compositions (structure). The null distribution of $H_{\text{non-SDEG},\, m}, m > 0$, is varied for each subtype, either when sample sizes are unequal across subtypes (**Eq. S5)** or when the non-SDEGs is unevenly distributed in the scatter plot. One example for the latter case is that $m$ subtypes are closer to each other than any others, causing these $m$ subtypes to have a larger condition probability of $H_{\text{non-SDEG},\, K-m}$.

#### By-products of aggregating genewise posterior probability

##### Estimating condition probability of one subtype being upregulated under the the null

More subtypes will increase the complexity of the null hypotheses. $P\{H_{\text{non-SDEG},\, m} | H_{\text{non-SDEG}}\}$, $m = 0, \dots, K-2$, only reflects the component weights of $(K-1)$ the null hypothesis types, identifying no specific subtype. By aggregating genewise posterior probability, we can obtain the probability of one subtype being upregulated conditioned on $H_{\text{non-SDEG}}$, which will affect the number of False Positive SDEGs of this subtype.

Suppose $H_{\text{non-SDEG},\, m}, m = 0, \dots, K-2$, has an equal prior probability for $(K-m)$ directions. Under the composition of multiple the null hypotheses, we get an estimated probability of one subtype being upregulated under the null:

$$\Pr\{up\ in\ k | H_{\text{non-SDEG}}\} = \frac{\sum_{m=0}^{K-2} \sum_{j=1}^{J} w_{\text{non-SDEG},\, m}(j) I\big(reOrder_{kj} \le K-m\big)/(K-m)}{\sum_{m=0}^{K-2} \sum_{j=1}^{J} w_{\text{non-SDEG},\, m}(j)}, \qquad (S9)$$

where $reOrder_{kj}$ is the reversed order of subtype $k$ sorted by expressions of gene $j$ across all subtypes. $I(*)$ indicates whether, for gene $j$, subtype $k$ is among the highest expressed $(K-m)$ subtypes and thus being upregulated with probability of $1/(K-m)$.

##### Landscape of gene expression patterns

There are $(2^K-1)$ types of gene expression patterns in a real dataset: genes exclusively expressed in $1, 2, \ldots$, or $K$ of subtypes. Genes unexpressed in any of subtypes have been eliminated during pre-processing. Aggregating the genewise posterior probability of the null hypotheses, $w_{\text{non-SDEG}, m}$, and of alternative hypotheses, $w_{\text{SDEG}}(j) = 1 - \sum_{n=0}^{K-2} w_{\text{non-SDEG}, n}(j)$, can present us a landscape of complex expression patterns as

$$P\{H_{\text{non-SDEG(A)}, m}\} = \frac{1}{J}\sum_{j=1}^{J} w_{\text{non-SDEG}, m}(j) I\left(\{k | reOrder_{kj} \leq K-m\} = A\right), \\ A \subseteq \{1, \ldots, K\}, |A| = K-m, 0 \leq m \leq K-2, \qquad (S10a)$$

$$P\{H_{\text{SDEG}(k)}\} = \frac{1}{J}\sum_{j=1}^{J} w_{\text{SDEG}}(j) I\left(reOrder_{kj} = 1\right), 1 \leq k \leq K, \qquad (S10b)$$

where $reOrder_{kj}$ is the reversed order of subtype $k$ sorted by expressions of gene $j$ across all subtypes. $A$ could be any subset of $\{1, \ldots, K\}$ with $(K-m)$ elements. $\{k | order_{kj} \leq K-m\}$ is a subset with elements being the indexes of $(K-m)$ highest expressed subtypes for gene $j$.

#### Within-subtype variance estimator

OVE-sFC test adopts an empirical Bayes moderated variance estimator used in "limma" that leverages information across genes by assuming a conjugate prior distribution $\sigma^2(j) \sim \nu_0 \sigma_0^2 / \mathcal{X}_{\nu_0}^2$[6]. As $t\text{-}stat_{k,l}(j)$ with moderated variance estimator $\tilde{\sigma}_j^2$ follows a t-distribution on $\nu_0 + N - K$ degrees of freedom, the increased degrees of freedom $\nu_0$ reflect the greater reliability associated with the smoothed variances. If mean-variance relationship exists in expression data, e.g. RNAseq data, "limma-voom" weights are incorporated into the linear modelling procedures to stabilize variance[7]. Other variance estimators designed for two-group differential analysis can also be easily modified and integrated into the multiple-group comparisons by OVE-sFC test. For example,

ROTS [8] adds a constant to the pooled variance estimator to optimize reproducibility across bootstrap resamplings.

##### Relevant peer SDEG selection methods

Two OVR methods, OVR-FC and OVR t-test, are included in peer method comparison. Note the degrees of freedom in OVR t-test (Welch's t-test) vary across genes, generating different the null distributions for each gene's test. Therefore, ranking genes based on OVR t-stat is different from that based on OVR t-test p-values.

We only calculate OVR-FC, OVR t-stat, and OVR t-test p-value for the highest expressed subtype. We do the same for OVE-FC, OVE-sFC (**Eq.1**), and OVE-sFC test p-value (**Eq.9**) and subtype-specific t-test p-value (**Eq.S8**). We use a classic method to calculate p-value for OVO t-test based on pairwise differential test[9]:

$$p\text{-}value = \max_{l \neq (K)} \{ p_{(K)l} \} \tag{S11}$$

where $p_{(K)l}$ is the p-value of differential test between subtype $(K)$ and subtype $l$. For a fair comparison between OVO t-test and OVE-sFC test, moderated variance estimator "limma" is applied to pairwise differential analyses in OVO t-test. As moderated variance estimator depends on the information across all genes for each subtype pair, all the $K(K-1)/2$ subtype pairs for all genes have to be tested. While OVO t-test leverages information from each subtype pair for variance modeling, OVE-sFC test leverages information from all subtypes. Moreover, OVE-sFC test re-estimates p-value by novel permutation scheme to model the complex the null distribution.

#### Simulations and Evaluations

##### Simulation study for validating OVE-sFC test statistics on type I error

We simulated a set of microarray experiments with 10000 genes,where baseline expression levels are sampled from real microarray data for the purified replicates in GSE19380 [10]. A portion of these genes are modeled as housekeeping genes under $H_{\text{non-SDEG},0}$ that keep their baseline expression values in all subtypes. The remaining genes are adjusted to have the same-level expression upregulation in at least two subtypes, mimicking other kinds of the null hypotheses. The upregulation levels are drawn from a uniform distribution in scatter space (black dash line in **Fig. S1a**). Genewise variation was generated from an inverse chi-square distribution, $\sigma^2(j) \sim \nu_0 \sigma_0^2 / \mathcal{X}_{\nu_0}^2$, with the prior degree of freedom $\nu_0$ being 5 or 40 related to less or more stabilized variances, respectively. $\sigma_0$ is set to be 0.2, 0.5, or 0.8 to check the performance under different noisy scenarios. **Fig. 1a** and **Fig. S2** show type I error control in the setting of three subtypes with balanced/unbalanced sample sizes. While permuting all subtypes generates a less dispersed the null distribution, and permuting only the top two subtypes results in a more compact distribution, our posterior weighted permutation scheme can achieve an automatic balance. Moreover, when the percentage of housekeeping genes (i.e. $H_{\text{non-SDEG, 0}}$) increases, the the null distribution of OVE-sFC test statistics tends to be that generated from permuting all three subtypes. Conversely, those calculated by permuting the top two subtypes approximate true p-values in $H_{\text{non-SDEG, 1}}$-dominated experiments. Comparison of results with different $\sigma_0$ demonstrates our test can control the type I error rate even in very noisy scenarios. However, small $\nu_0$ generates a modestly inflated type I error control, especially when sample sizes are small and the true $H_{\text{non-SDEG, 1}}$ dominates. This effect may be due to a less reliable estimation of the moderated variance caused by small $\nu_0$ and sample sizes.

The estimated conditional probabilities of $H_{\text{non-SDEG, 0}}$, $P\{H_{\text{non-SDEG, 0}} | H_{\text{non-SDEG}}\}$ in our model are expected to match the true proportions of housekeeping genes. As seen in **Fig. S3**, in noisy scenarios with high $\sigma_0$, $P\{H_{\text{non-SDEG, 0}} | H_{\text{non-SDEG}}\}$ tends to get over-estimated, implying that many genes sampled from the true $H_{\text{non-SDEG, 1}}$ are treated as being from $H_{\text{non-SDEG, 0}}$ in certain weights. In fact, these true $H_{\text{non-SDEG, 1}}$ genes have no significantly large $\Delta_1(j)$, which is the difference between the top two subtypes and the third subtype. Thus, the the null distributions generated from them are expected to be intermediate between those from true $H_{\text{non-SDEG, 0}}$ (permuting three subtypes) and from true $H'_{\text{non-SDEG, 1}}$ (permuting top 2 subtypes).

Subtype-specific OVE-sFC test conducts the tests for SDEGs of each subtype separately, having the benefit that all subtypes a broadly similar type I error rate (**Fig. 1a**). Otherwise, the subtype consisting of a smaller sample size will contribute more False Positive SDEGs. Note that subtype-specific p-values could be larger than 1 after multiple-tail scaling (**Fig. 1b**) and need to be truncated at 1. While we only calculate the p-value for the subtype with the largest group mean to reduce the computational burden, we ignore the possibility that genes with large p-values (around one) could have smaller p-values associated with another subtype. Since these genes are not likely to be valid SDEGs, it is not necessary to provide their precise p-values.

All the above analyses are repeated for SDEG identification involving five subtypes, producing p-values that control error rates correctly over a wide range of simulation scenarios (**Fig. 2a**). Since we randomly assigned genes under $H_{\text{non-SDEG},\,m}, m > 0$ without considering subtypes in the simulations above, the the null hypothesis compositions are almost the same for all subtypes. To check the type I error rate under unbalanced the null hypothesis compositions, five baseline profiles are generated by simulating two cell lines but assigning one to two subtypes and the other to three subtypes, so that the first cell line's up-regulated genes become two subtypes' $H_{\text{non-SDEG},\,3}$ genes, while the second cell line's up-regulated genes become three subtypes' $H_{\text{non-SDEG},\,2}$ genes. No true SDEGs exist for any of the five subtypes. OVE-sFC test (overall or subtype-specific) controls type I error rates well but with more False Positive SDEGs in the first two subtypes (**Fig. 2b**). The unbalance of the null hypotheses leads to different probability estimates of a specific subtype being upregulated (**Eq. S9**). As simulated up-regulated genes in two cell lines are the same but are divided into two or three subtypes evenly, the first two subtypes have a higher probability of being upregulated, thus more False Positive SDEGs are expected (**Fig. S4**). In our simulations, when the ratio of housekeeping genes is high, the unbalance becomes slight and thus approximately 1/5 of False Positive SDEGs are allocated to each subtype. Conversely, a low ratio of housekeeping genes intensifies the uneven distribution of non-SDEGs in the scatter plot, increasing the number of False Positive SDEGs detected in the first two subtypes. Subtype-specific tests can reduce this impact.

#### Simulation study for assessing the power of OVE-sFC test statistics

A well-designed test can maximize power while controlling FDR below the expected level. In data sets involving true SDEGs, we evaluated several SDEG selection methods with respect to FDR control and pAUC. We studied whether the tests can control the FDR well by checking whether the true FDR with q-value (estimated by 'fdrtool' package [3]) at 0.05 cutoff is also around 0.05. The area under the receiver operating characteristic (ROC) curve reflects whether the methods are able to rank true SDEGs above true non-SDEGs. In practical analyses, we emphasize the sensitivity or power of SDEG detection methods when the False Positive Rate (FPR) is significantly low, such as 0.05/0.01 cutoff. Hence, the partial area under curve (pAUC) with specificity larger than 0.95/0.99 (equivalent to FPR less than 0.05/0.01) was used to evaluate the power of each method. Another reason here is that the emergence of mismatched detections (True Positive but associated with incorrect subtypes) with low specificity may exhibit a false high power.

Simulation settings are similar to the above subsection except that a portion of genes are designated as SDEGs with upregulation in one of the subtypes. We conducted three simulation sets to assess the power of OVE-FC/sFC test statistics and peer methods: (1) microarray simulations with variance drawn from an inverse chi-square distribution ($K$=3); (2) microarray simulations with variance sampled from real data ($K$=7); (3) RNASeq simulations with variance sampled from real data ($K$=7). The latter two simulation sets are more challenging with more subtypes and noisy RNASeq data, which help show the benefit of using all subtypes for modeling by the proposed OVE-sFC test.

In the first simulation set, genewise variation was drawn from $\sigma^2(j) \sim \frac{\nu_0 \sigma_0^2}{\chi_{\nu_0}^2}$, with the prior degree of freedom $\nu_0$ being 5 or 40 related to less or more stabilized variances, respectively. $\sigma_0$ is set to be 0.2, 0.5, or 0.8 to check the performance under different noisy scenarios. 20% of genes are upregulated in one subtype with fold change following a uniform distribution in the Ternary plot (**Fig. S1a**), among which fold change ranges from 1 to $+\infty$. The SDEGs with small fold change (close to 1) are non-ideal SDEGs. 40% of the remaining genes are under $H_{\text{non-SDEG},\,0}$ and 40% under $H_{\text{non-SDEG},\,1}$. Two scenarios were tested with unbalanced sample sizes or unbalanced the null hypothesis compositions. In the former one, sample size in each subtype is 3, 6, and 9, respectively, while $H_{\text{non-SDEG},\,1}$ genes are distributed evenly. In the latter one, each subtype has three samples and $H_{\text{non-SDEG},\,1}$ genes only appear in the first two subtypes, which is a common case in real data: two subtypes are closer to each other than to the other subtypes. Each simulation was repeated 20 times. Both the overall and subtype-specific OVE-sFC tests could control FDR around the expected 0.05 level at q-value cutoff 0.05, with modestly weaker control in the case of small $\nu_0$ (**Fig. S5**). OVR t-test is too liberal, while OVO t-test is conservative. Furthermore, a subtype-specific test can alleviate the unbalance of False Positive SDEGs among subtypes. Other peer methods detected more False Positive SDEGs in subtypes with a small sample size or with a large probability of being upregulated under $H_{\text{non-SDEG}}$ (**Eq. S9**). In terms of pAUC, OVE-sFC, OVE-FC and OVO t-test test achieved better performance than OVR methods (**Table S1**). OVE-sFC outperforms OVE-FC due to many non-ideal SDEGs in this simulation set.

In the second and third simulation set, genewise variation was sampled from microarray data GSE28490 or RNAseq data GSE60424. To keep potential mean-variance trend, we divided genes from real data into 100 buckets based on their logarithmic expression means and randomly selected a real variance for each simulated gene from the bucket which the simulated expression mean fell in. 100 SDEGs were simulated for each subtype while the remaining genes were simulated as different yet realistic types of non-SDEGs, $H_{\text{non-SDEG},m}, m = 0,1,\ldots,5$. The the null hypothesis structure, i.e. the percentage of different types of non-SDEGs for each subtype, was generated based on the estimated probability of gene expression patterns (**Eq. S10a**) from real data. Since this the null hypothesis structure from real data is somewhat unbalanced, we also simulated a balanced the null hypothesis structure with each subtype having the same percentage of different types of non-SDEGs and sample size being balanced or unbalanced. To compare the performance of detecting ideal/strict SDEGs (significantly large fold change) or detecting more realistic SDEGs (sufficiently large fold change), the fold changes of simulated SDEGs were drawn from a uniform distribution with different range: [2,20], [5,20], or [10,20]. Both the overall and subtype-specific OVE-sFC tests could control FDR around the expected 0.05 level at q-value cutoff 0.05, with slightly weaker control for noisy RNASeq data with a small sample size (**Fig. S6**). In terms of pAUC, OVE-sFC tests and OVE-FC approach the highest power in detecting true SDEGs (**Table S2-S3**, **Fig. 3**, **and Fig. S7-S8**).

#### Real datasets to detect human immune cell markers

We used two real microarray datasets, GSE28490 (Roche) and GSE28491 (HUG), which contain mRNA expression profiles of seven immune cell types (B cells, CD4+ T cells, CD8+ T cells, NK cells, monocytes, neutrophils, and eosinophils) isolated from healthy human blood pools [11]. Two datasets used the same protocols for cell isolation and sample processing on two independent panels of donors. All cell types have five samples in each dataset except monocyte has ten in Roche dataset, with few samples (2 neutrophils in Roche, 1 eosinophil in Roche, 2 eosinophils in HUG) removed from datasets as outliers (**Table S7**). After eliminating low expressed probesets with an average $\log_2$ RMA signal value > 6 in none of the cell type groups, 12022/11339 probesets in Roche/ HUG datasets were considered to be expressed above background and thus available for use with OVE-sFC test.

#### Evaluate SDEGs by supervised deconvolution performance

##### Supervised deconvolution by CAM score

There are two major classes of supervised deconvolution approaches to quantify proportions of subtypes in mixtures: fitting coefficients for the linearly modeled relationship between mixture and pure expression levels; or scoring each subtype by expression levels of markers in mixtures[12]. The former approach estimates the absolute fraction of each subtype in a heterogeneous sample. The latter one provides relative scores that are comparable across samples but not comparable between subtypes. However, the second class has its own advantages. **(1)** It only assumes the indexes of markers are unchangeable, so that the markers selected based on pure expression levels can help deconvolute mixtures in different microenvironments, even when measured on other platforms. **(2)** Scores are computed for each subtype individually and thus less affected by each other and unknown subtypes. **(3)** Scores are able to achieve a higher correlation with ground truth proportions, since they focus on catching the dynamic trend across samples, not the absolute proportion values. Generally, the factors affecting coefficient fitting are complex, whereas scoring one subtype relies mostly on the precision of the indexes of selected markers. Therefore, we adopted a scoring approach for supervised deconvolution and use its performance to evaluate selected markers.  The details of the score estimation are as below.

The expression level of the $j$th gene in the $i$th mixture sample, $x_{ij}$, is modeled as a linear combination of the expression levels of that gene across the $K$ subtypes present in the mixture, weighted by their constituent proportions $a_{ik}$:

$$x_{ij} = \sum_{k=1}^{K} a_{ik} s_k(j)\,. \qquad (S12)$$

Consider a set of N heterogeneous samples and denote $x(j) = [x_{1j}, \dots, x_{Nj}], a_k = [a_{1k}, \dots, a_{Nk}]$. We can re-write **Eq. S12** as

$$x(j) = \sum_{k=1}^{K} s_k(j) a_k . \qquad (S13)$$

As molecular markers are exclusively expressed in only one of each subtype, we have

$$x(j_{MG\text{-}k}) \approx s_k(j_{MG\text{-}k}) a_k, \qquad (S14)$$

where $j_{MG\text{-}k}$ is the index of any marker of subtype $k$. Therefore, the expression levels of markers are proportional to the constituent fraction of certain subtypes. After applying a sum-to-one standardization to $x(j_{MG\text{-}k})$, their space median can be used as the score for subtype $k$:

$$\tilde{a}_k = SpaceMedian\{\tilde{x}(j_{MG\text{-}k})\}. \qquad (S15)$$

“l1median” function in R package “pcaPP”[13] is used to find space median.

Such score estimation approach is the same as part of an unsupervised deconvolution algorithm - “CAM” [14]. We can call **Eq. S15** a CAM score. There are also other score methods among which MCP-counter score [15], the geometric mean of $x(j_{MG\text{-}k})$, is the most similar one to CAM score. We expect the space median operation to enhance the robustness of scores. Sample-specific enrichment score, e.g. ssGSEA, has also been proposed to infer subtypes in tissues [16, 17], but its relationship with the true subtype proportion is non-linear and unclear.

##### In silico simulated mixtures

We generate gene expression readout of *in silico* heterogeneous samples following the linear mixing model in **Eq. S12** with additional biological variation and technical variation:

$$x_{ij} = \sum_{k=1}^{K} a_{ik}\left(s_k(j) + \Delta s_{ik}(j)\right) + \varepsilon_{ij}, \qquad (S16)$$

where $\Delta s_{ik}(j)$ reflects subtype-specific biological variation across samples and $\varepsilon_{ij}$ is technical noise of measurement. In the first and the third simulation (**Fig. 5a, 5c**), $s_k(j)$ is cell-type specific mRNA expression mean (averaged across samples in log2 scale and transformed to original scale) in GSE28491. $\Delta s_{ik}(j)$ and $\varepsilon_{ij}$ follows zero-mean Gaussian distribution, with variance drawn from the inverse chi-square distribution with $\sigma_0$ being 0.5 and 0.1, $\nu_0$ being 10 and 5, respectively. In the second simulation (**Fig. 5b**), as GSE60424 has almost 20 samples per cell type, one sample was randomly selected from each cell type and its RNAseq count profile could be treated as $s_k(j)$ with biological and technical variance already included. Mixing proportions, $a_{ik}, k = 1, \dots, K$, were drawn randomly from a flat Dirichlet distribution and used in the three simulations.

### Supplementary Tables

**Table S1** pAUC (FPR<0.05) obtained from Microarray simulations involving 3 subtypes and with various experimental settings

| | | $\nu 0 = 5$ | | | $\nu 0 = 40$ | | |
|---|---|---|---|---|---|---|---|
| | | $\sigma = 0.2$ | $\sigma = 0.5$ | $\sigma = 0.8$ | $\sigma = 0.2$ | $\sigma = 0.5$ | $\sigma = 0.8$ |
| Balanced The null Hypothesis Structure<br><br>Unbalanced Sample Size (n=3,6,9) | ANOVA | 0.53747 | 0.5402 | 0.54266 | 0.53794 | 0.54081 | 0.54344 |
| | OVR-FC | 0.67778 | 0.65644 | 0.62664 | 0.68155 | 0.66657 | 0.64723 |
| | OVR t-stat | 0.9093 | 0.77705 | 0.6904 | 0.92751 | 0.80715 | 0.71382 |
| | OVR t-test | 0.82649 | 0.70774 | 0.64513 | 0.84036 | 0.73137 | 0.66212 |
| | OVO t-test | 0.94982 | 0.8569 | 0.76983 | 0.96193 | 0.89067 | 0.80318 |
| | OVE-FC | 0.94292 | 0.84033 | 0.74264 | 0.96179 | 0.88991 | 0.80195 |
| | OVE-sFC | **0.95116** | **0.85907** | **0.77208** | **0.96196** | **0.89074** | **0.80403** |
| | sub OVE-sFC | **0.95119** | **0.85915** | **0.77301** | **0.96211** | **0.89078** | **0.80424** |
| Unbalanced The null Hypothesis Structure<br><br>(n=3,3,3) | ANOVA | 0.54128 | 0.54473 | 0.54533 | 0.54105 | 0.54472 | 0.54411 |
| | OVR-FC | 0.70822 | 0.67034 | 0.62272 | 0.71086 | 0.68668 | 0.65558 |
| | OVR t-stat | 0.92793 | 0.78802 | 0.68392 | 0.94068 | 0.81609 | 0.70628 |
| | OVR t-test | 0.89682 | 0.73482 | 0.6429 | 0.91385 | 0.75827 | 0.65584 |
| | OVO t-test | 0.93685 | 0.8218 | 0.72267 | 0.95218 | 0.86223 | 0.76863 |
| | OVE-FC | 0.93113 | 0.80302 | 0.69664 | 0.9522 | 0.86167 | 0.7676 |
| | OVE-sFC | **0.93887** | **0.82354** | **0.72451** | **0.95211** | **0.86229** | **0.76811** |
| | sub OVE-sFC | **0.94075** | **0.82749** | **0.72821** | **0.95383** | **0.86651** | **0.77229** |

**Table S2** pAUC (FPR<0.05 and 0.01) obtained from Microarray simulations involving 7 subtypes and with various experimental settings

| | | pAUC (FPR<0.05) | | | pAUC (FPR<0.01) | | |
|---|---|---|---|---|---|---|---|
| | | FC ∈ [2,20] | FC ∈ [5,20] | FC ∈ [10,20] | FC ∈ [2,20] | FC ∈ [5,20] | FC ∈ [10,20] |
| Unbalanced The null Hypothesis Structure 3 samples /per subtype | ANOVA | NA | NA | NA | NA | NA | NA |
| | OVR-FC | NA | 0.50997 | 0.56310 | NA | NA | 0.50035 |
| | OVR t-stat | 0.81362 | 0.92408 | 0.96226 | 0.80365 | 0.91644 | 0.95595 |
| | OVR t-test | 0.55572 | 0.57203 | 0.59205 | 0.56113 | 0.57531 | 0.59397 |
| | OVO t-test | 0.93403 | 0.98283 | 0.98955 | 0.89324 | 0.96797 | 0.97626 |
| | OVE-FC | 0.93612 | **0.99533** | **0.99878** | 0.81569 | **0.98423** | **0.99759** |
| | OVE-sFC | **0.94797** | 0.98833 | 0.99348 | **0.92393** | 0.97953 | 0.98765 |
| | sub OVE-sFC | **0.95052** | 0.98897 | 0.99384 | **0.92372** | 0.98028 | 0.98764 |
| Balanced The null Hypothesis Structure 3 samples /per subtype | ANOVA | NA | NA | NA | NA | NA | NA |
| | OVR-FC | NA | 0.50671 | 0.55154 | NA | NA | 0.50275 |
| | OVR t-stat | 0.82253 | 0.92019 | 0.94936 | 0.80918 | 0.91560 | 0.94403 |
| | OVR t-test | 0.56269 | 0.58333 | 0.58408 | 0.56490 | 0.58558 | 0.58595 |
| | OVO t-test | 0.92870 | 0.98165 | 0.98961 | 0.88065 | 0.96383 | 0.97822 |
| | OVE-FC | 0.93763 | **0.99568** | **0.99770** | 0.82500 | **0.98386** | **0.98897** |
| | OVE-sFC | **0.95044** | 0.98929 | 0.99317 | **0.91775** | 0.98127 | **0.98948** |
| | sub OVE-sFC | **0.95062** | 0.98935 | 0.99316 | **0.91830** | 0.98163 | **0.98947** |
| Balanced The null Hypothesis Structure Unbalanced Sample Size (3,3,3,4,5,5,5) | ANOVA | NA | NA | NA | NA | NA | NA |
| | OVR-FC | NA | 0.50686 | 0.54375 | NA | NA | 0.50004 |
| | OVR t-stat | 0.79462 | 0.90719 | 0.94392 | 0.78328 | 0.89854 | 0.93593 |
| | OVR t-test | 0.52553 | 0.54197 | 0.55172 | 0.52574 | 0.54076 | 0.55029 |
| | OVO t-test | 0.95338 | 0.99001 | 0.99563 | 0.92252 | 0.98104 | 0.98687 |
| | OVE-FC | 0.95395 | **0.99763** | **0.99882** | 0.85616 | **0.98841** | **0.99423** |
| | OVE-sFC test | **0.96077** | 0.99411 | 0.99711 | **0.94228** | 0.98681 | 0.99301 |
| | sub OVE-sFC | **0.96101** | 0.99418 | 0.99711 | **0.94222** | 0.98693 | 0.99317 |

**Table S3** pAUC (FPR<0.05 and 0.01) obtained from RNAseq simulations involving 7 subtypes and with various experimental settings

| | | pAUC (FPR<0.05) | | | pAUC (FPR<0.01) | | |
|---|---|---|---|---|---|---|---|
| | | $FC \in [2,20]$ | $FC \in [5,20]$ | $FC \in [10,20]$ | $FC \in [2,20]$ | $FC \in [5,20]$ | $FC \in [10,20]$ |
| Unbalanced The null Hypothesis Structure<br><br>3 samples /per subtype | ANOVA | NA | NA | NA | NA | NA | NA |
| | OVR-FC | 0.50007 | 0.51649 | 0.59034 | 0.50120 | 0.50260 | 0.52331 |
| | OVR t-stat | 0.63528 | 0.76419 | 0.81669 | 0.60403 | 0.70409 | 0.77145 |
| | OVR t-test | 0.54461 | 0.60287 | 0.63803 | 0.53923 | 0.58136 | 0.62174 |
| | OVO t-test | 0.77600 | 0.88615 | 0.93747 | 0.68080 | 0.76939 | 0.85403 |
| | OVE-FC | 0.75622 | **0.93221** | **0.97183** | 0.62926 | 0.80988 | **0.90791** |
| | OVE-sFC | **0.79344** | 0.91728 | 0.96178 | **0.68795** | **0.81828** | 0.89212 |
| | sub OVE-sFC | **0.79869** | 0.91810 | 0.96212 | **0.69353** | **0.81877** | 0.89454 |
| Unbalanced The null Hypotheses<br><br>20 samples /per subtype | ANOVA | NA | NA | NA | NA | NA | NA |
| | OVR-FC | NA | 0.53594 | 0.62450 | NA | 0.50883 | 0.53201 |
| | OVR t-stat | 0.61767 | 0.75120 | 0.82295 | 0.60970 | 0.72330 | 0.80238 |
| | OVR t-test | 0.50731 | 0.55117 | 0.60501 | 0.51059 | 0.54524 | 0.59214 |
| | OVO t-test | 0.97075 | 0.99193 | 0.99578 | **0.96192** | 0.98840 | 0.99354 |
| | OVE-FC | **0.97611** | 0.99369 | 0.99660 | 0.95103 | 0.98974 | 0.99263 |
| | OVE-sFC | 0.97232 | **0.99448** | **0.99782** | 0.95100 | **0.99131** | **0.99632** |
| | sub OVE-sFC | 0.97363 | **0.99438** | **0.99770** | 0.95323 | **0.99093** | **0.99645** |
| Balanced The null Hypothesis Structure<br><br>3 samples /per subtype | ANOVA | NA | NA | NA | NA | NA | NA |
| | OVR-FC | NA | 0.52764 | 0.59584 | NA | 0.50463 | 0.52588 |
| | OVR t-stat | 0.63400 | 0.76959 | 0.82773 | 0.60631 | 0.72285 | 0.77652 |
| | OVR t-test | 0.54489 | 0.61288 | 0.63604 | 0.54151 | 0.59807 | 0.62051 |
| | OVO t-test | 0.77225 | 0.89831 | 0.93346 | 0.67555 | 0.79982 | 0.86136 |
| | OVE-FC | 0.74739 | **0.92323** | **0.97642** | 0.61980 | 0.78167 | **0.91866** |
| | OVE-sFC | **0.77921** | 0.91873 | 0.96874 | **0.68426** | **0.81631** | 0.91196 |
| | sub OVE-sFC | **0.77931** | 0.91857 | 0.96867 | **0.68397** | **0.81483** | 0.91164 |
| Balanced The null Hypothesis Structure<br><br>Unbalanced Sample Size (3,3,3,4,5,5,5) | ANOVA | NA | NA | NA | NA | NA | NA |
| | OVR-FC | NA | 0.51603 | 0.57669 | NA | 0.50377 | 0.51511 |
| | OVR t-stat | 0.61345 | 0.74834 | 0.79636 | 0.59512 | 0.70675 | 0.76168 |
| | OVR t-test | 0.53701 | 0.58239 | 0.62472 | 0.53539 | 0.56349 | 0.60675 |
| | OVO t-test | 0.79078 | 0.91174 | 0.93552 | 0.69959 | 0.82953 | 0.87458 |
| | OVE-FC | 0.75689 | **0.95120** | **0.97319** | 0.61997 | **0.85831** | **0.92000** |
| | OVE-sFC | **0.79951** | 0.92714 | 0.95949 | **0.70584** | 0.84436 | 0.89120 |
| | sub OVE-sFC | **0.79978** | 0.92689 | 0.95925 | **0.70544** | 0.84413 | 0.88982 |

**Table S4** Counts of cell-type specific markers under certain threshold

| Threshold | Subtype | Measured only in Roche | Markers only in Roche | Markers in both | Markers only in HUG | Measured only in HUG | Total |
|---|---|---|---|---|---|---|---|
| q-value <0.05 | B cells | 70 | 266 | 474 | 528 | 39 | 1377 |
| | CD4+ T cells | 6 | 44 | 28 | 54 | 1 | 133 |
| | CD8+ T cells | 8 | 55 | 7 | 33 | 2 | 105 |
| | NK cells | 83 | 563 | 208 | 68 | 13 | 935 |
| | Eosinophils | 37 | 301 | 204 | 106 | 11 | 659 |
| | Monocytes | 51 | 475 | 630 | 463 | 55 | 1674 |
| | Neutrophils | 43 | 519 | 626 | 256 | 76 | 1520 |
| q-value <0.001 | B cells | 56 | 209 | 264 | 102 | 13 | 644 |
| | CD4+ T cells | 2 | 12 | 4 | 4 | 0 | 22 |
| | CD8+ T cells | 3 | 20 | 3 | 1 | 0 | 27 |
| | NK cells | 52 | 255 | 85 | 8 | 4 | 404 |
| | Eosinophils | 27 | 208 | 55 | 13 | 4 | 307 |
| | Monocytes | 46 | 427 | 260 | 67 | 18 | 818 |
| | Neutrophils | 24 | 452 | 173 | 31 | 32 | 712 |
| Corrected p-value <0.05 | B cells | 44 | 134 | 181 | 55 | 9 | 423 |
| | CD4+ T cells | 1 | 4 | 2 | 0 | 0 | 7 |
| | CD8+ T cells | 0 | 14 | 1 | 0 | 0 | 15 |
| | NK cells | 40 | 185 | 37 | 1 | 3 | 266 |
| | Eosinophils | 19 | 49 | 24 | 15 | 0 | 107 |
| | Monocytes | 36 | 289 | 137 | 20 | 5 | 487 |
| | Neutrophils | 13 | 203 | 56 | 13 | 15 | 300 |
| Corrected p-value <0.001 | B cells | 41 | 116 | 148 | 44 | 9 | 358 |
| | CD4+ T cells | 1 | 5 | 1 | 0 | 0 | 7 |
| | CD8+ T cells | 0 | 13 | 1 | 0 | 0 | 14 |
| | NK cells | 32 | 158 | 18 | 2 | 3 | 213 |
| | Eosinophils | 19 | 61 | 12 | 2 | 0 | 94 |
| | Monocytes | 33 | 296 | 60 | 1 | 0 | 390 |
| | Neutrophils | 13 | 255 | 3 | 1 | 3 | 275 |

**Table S5** Statistics of CD4+ T cell markers detected in both Roche and HUG (q<0.05)

| | Probe /Probeset | Gene Symbol | Roche | | | HUG | | |
|---|---|---|---|---|---|---|---|---|
| | | | OVE-FC | p-value | q-value | OVE-FC | p-value | q-value |
| 1 | 206492_at | FHIT | 3.302 | 3.89E-17 | 8.87E-16 | 4.42 | 2.08E-14 | 7.36E-13 |
| 2 | 229070_at | ADTRP | 7.068 | 4.25E-13 | 4.60E-12 | 7.365 | 3.21E-06 | 3.02E-05 |
| 3 | 209442_x_at | ANK3 | 2.397 | 6.33E-13 | 6.48E-12 | 1.747 | 2.79E-04 | 1.21E-03 |
| 4 | 206385_s_at | ANK3 | 3.227 | 3.90E-10 | 2.71E-09 | 2.696 | 2.54E-04 | 1.13E-03 |
| 5 | 236341_at | CTLA4 | 2.919 | 4.88E-06 | 2.10E-05 | 3.233 | 1.65E-04 | 7.95E-04 |
| 6 | 1562731_s_at | MDS2 | 1.728 | 1.03E-05 | 4.12E-05 | 1.688 | 7.43E-04 | 2.61E-03 |
| 7 | 203410_at | AP3M2 | 2.061 | 1.91E-05 | 7.11E-05 | 3.621 | 4.33E-05 | 2.69E-04 |
| 8 | 1729_at | TRADD | 1.376 | 1.15E-04 | 3.35E-04 | 1.409 | 4.63E-03 | 1.07E-02 |
| 9 | 227641_at | FBXL16 | 1.388 | 4.61E-04 | 1.14E-03 | 1.911 | 1.07E-04 | 5.64E-04 |
| 10 | 217147_s_at | TRAT1 | 1.842 | 6.26E-04 | 1.48E-03 | 1.73 | 7.75E-03 | 1.59E-02 |
| 11 | 213135_at | TIAM1 | 1.47 | 7.25E-04 | 1.68E-03 | 1.798 | 2.19E-03 | 5.98E-03 |
| 12 | 203386_at | TBC1D4 | 1.398 | 8.25E-04 | 1.88E-03 | 2.034 | 2.06E-03 | 5.71E-03 |
| 13 | 227580_s_at | TECPR1 | 1.364 | 9.64E-04 | 2.16E-03 | 1.512 | 9.29E-03 | 1.82E-02 |
| 14 | 204773_at | IL11RA | 1.416 | 1.12E-03 | 2.46E-03 | 1.455 | 7.31E-03 | 1.52E-02 |
| 15 | 204777_s_at | MAL | 1.541 | 1.39E-03 | 2.98E-03 | 2.496 | 5.44E-05 | 3.22E-04 |
| 16 | 1557733_a_at | CHRM3-AS2 | 1.643 | 1.60E-03 | 3.38E-03 | 3.211 | 5.12E-05 | 3.06E-04 |
| 17 | 40016_g_at | MAST4 | 1.351 | 4.64E-03 | 8.55E-03 | 1.679 | 1.68E-03 | 4.88E-03 |
| 18 | 225613_at | MAST4 | 1.524 | 4.88E-03 | 8.94E-03 | 2.116 | 2.22E-03 | 6.04E-03 |
| 19 | 224832_at | DUSP16 | 1.274 | 8.84E-03 | 1.51E-02 | 1.712 | 8.09E-03 | 1.64E-02 |
| 20 | 213028_at | NFRKB | 1.171 | 8.85E-03 | 1.51E-02 | 1.243 | 3.44E-02 | 4.74E-02 |
| 21 | 203717_at | DPP4 | 1.394 | 1.46E-02 | 2.35E-02 | 1.834 | 4.14E-04 | 1.66E-03 |
| 22 | 225611_at | MAST4 | 1.366 | 1.88E-02 | 2.93E-02 | 2.131 | 1.33E-04 | 6.64E-04 |
| 23 | 206545_at | CD28 | 1.307 | 2.16E-02 | 3.28E-02 | 1.644 | 2.00E-02 | 3.22E-02 |
| 24 | 56197_at | PLSCR3 | 1.176 | 2.16E-02 | 3.29E-02 | 1.332 | 8.65E-03 | 1.73E-02 |
| 25 | 210439_at | ICOS | 1.338 | 2.29E-02 | 3.45E-02 | 2.262 | 1.31E-03 | 3.99E-03 |
| 26 | 220048_at | EDAR | 1.289 | 2.84E-02 | 4.15E-02 | 1.565 | 2.24E-03 | 6.07E-03 |
| 27 | 211675_s_at | MDFIC | 1.378 | 2.88E-02 | 4.20E-02 | 1.663 | 5.97E-03 | 1.30E-02 |
| 28 | 225478_at | MFHAS1 | 1.327 | 3.23E-02 | 4.63E-02 | 1.96 | 1.22E-03 | 3.78E-03 |

**Table S6** Statistics of CD8+ T cell markers detected in both Roche and HUG (q<0.05)

| | Probe /Probeset | Gene Symbol | Roche | | | HUG | | |
|---|---|---|---|---|---|---|---|---|
| | | | OVE-FC | p-value | q-value | OVE-FC | p-value | q-value |
| 1 | 215332_s_at | CD8B | 38.888 | 1.15E-31 | 8.90E-30 | 19.368 | 1.54E-20 | 2.77E-18 |
| 2 | 205758_at | CD8A | 5.635 | 4.81E-13 | 5.12E-12 | 3.485 | 7.02E-05 | 3.99E-04 |
| 3 | 203413_at | NELL2 | 2.741 | 7.08E-09 | 4.02E-08 | 1.898 | 9.87E-04 | 3.23E-03 |
| 4 | 206666_at | GZMK | 3.963 | 4.18E-07 | 2.04E-06 | 4.644 | 1.12E-05 | 8.73E-05 |
| 5 | 209871_s_at | APBA2 | 1.815 | 8.94E-05 | 2.70E-04 | 1.575 | 9.05E-03 | 1.79E-02 |
| 6 | 226474_at | NLRC5 | 1.497 | 3.16E-03 | 6.13E-03 | 1.439 | 2.02E-02 | 3.24E-02 |
| 7 | 225803_at | FBXO32 | 1.736 | 1.52E-02 | 2.43E-02 | 2.066 | 1.14E-03 | 3.60E-03 |

**Table S7** Sample size of each cell type in four datasets

| | GSE28490 | GSE28491 | GSE60424 | GSE72056* |
|---|---|---|---|---|
| B cells | 5 | 5 | 20 | 628 |
| CD4+ T cells | 5 | 5 | 20 | 873 |
| CD8+ T cells | 5 | 5 | 20 | 1099 |
| Regulatory T cells | NA | NA | NA | 141 |
| NK cells | 5 | 5 | 14 | 89 |
| Eosinophils | 4 | 3 | NA | NA |
| Macrophages/Monocytes | 10 | 5 | 20 | 170 |
| Neutrophils | 3 | 5 | 20 | NA |
| Plasmacytoid dendritic cells | 5 | NA | NA | 26 |
| Myeloid dendritic cells | 5 | NA | NA | NA |
| Endothelial cells | NA | NA | NA | 71 |
| Cancer associated fibroblasts | NA | NA | NA | 96 |

*Cell type labels are from single-cell classification results [18]

### Supplementary Figure

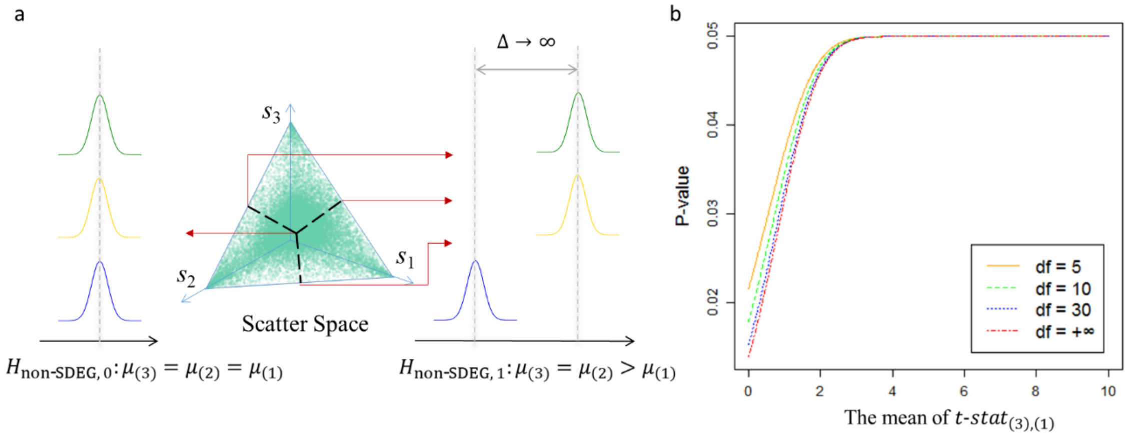


**Figure S1.** Two extreme cases of non-SDEGs when the subtype number is three. Fixing the critical value to be that of two-group t-test p = 0.05, p-values approach the upper bound when two subtypes are drawn from the same population with much larger expressions than the third. On the contrary, p-values decrease to be minimum when all three subtypes are drawn from the same population.

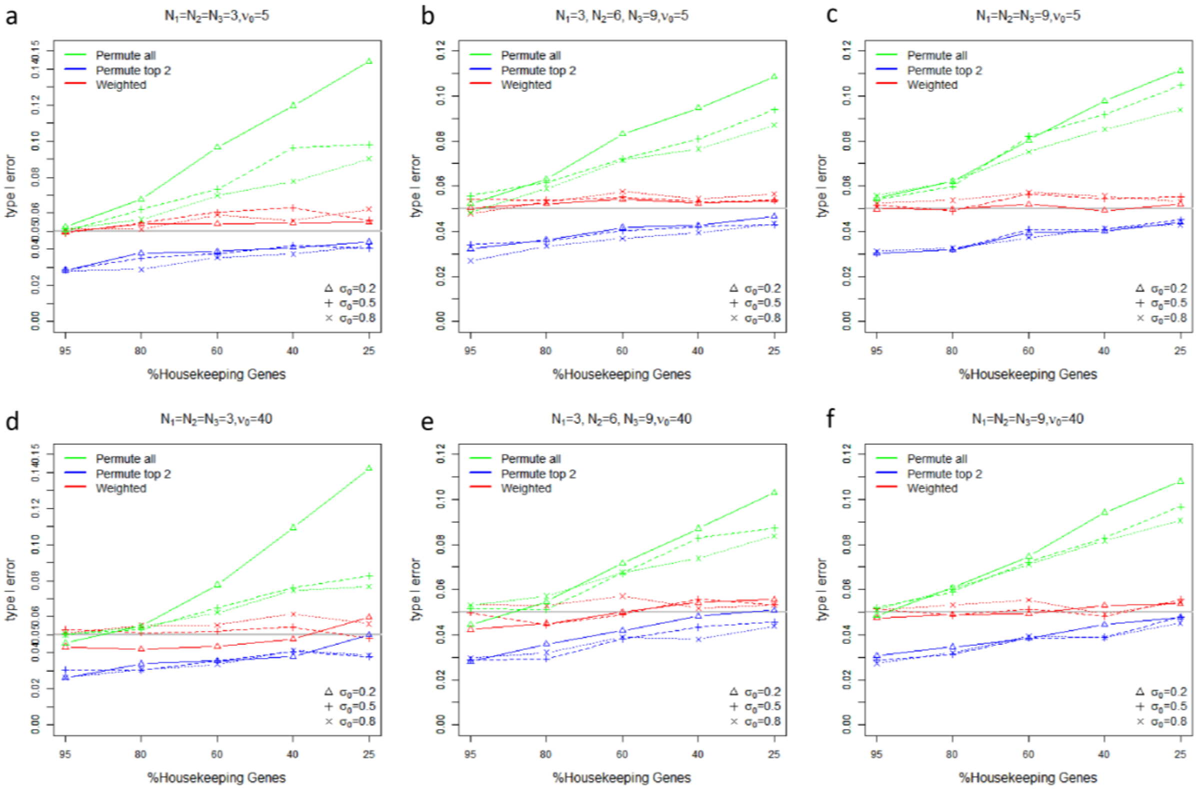


**Figure S2.** Comparisons of type I error rates from different permutation schemes under different settings of noisy scenarios and sample sizes (housekeeping genes: 95%, 80%, 60%, 40%, or 25%; $\sigma_0$: 0.2, 0.5, or 0.8; $\nu_0$: 5(top), or 40(bottom)).

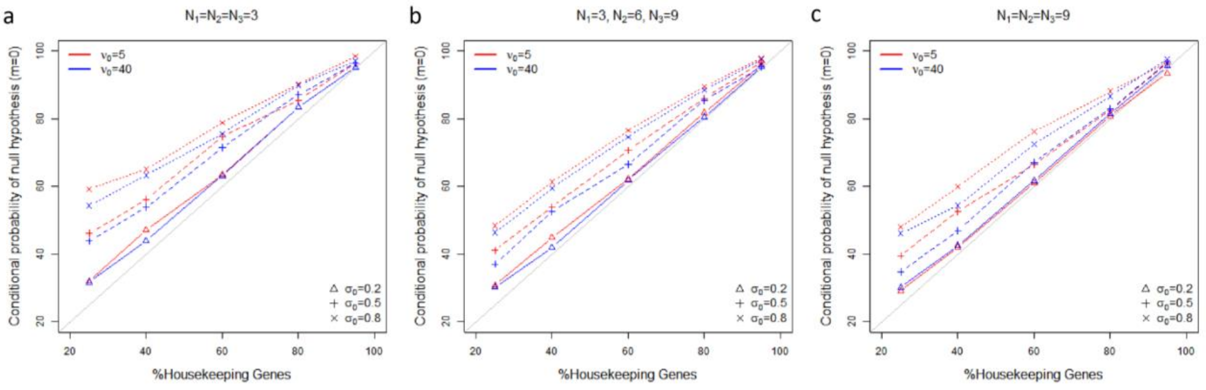


**Figure S3**. Comparisons of estimated conditional probabilities of the the null hypothesis $H_{\text{non-SDEG, 0}}$, $P\left\{H_{\text{non-SDEG, 0}} \middle| H_{\text{non-SDEG}}\right\}$, versus the true proportions of housekeeping genes, under different settings of noisy scenarios and sample sizes.

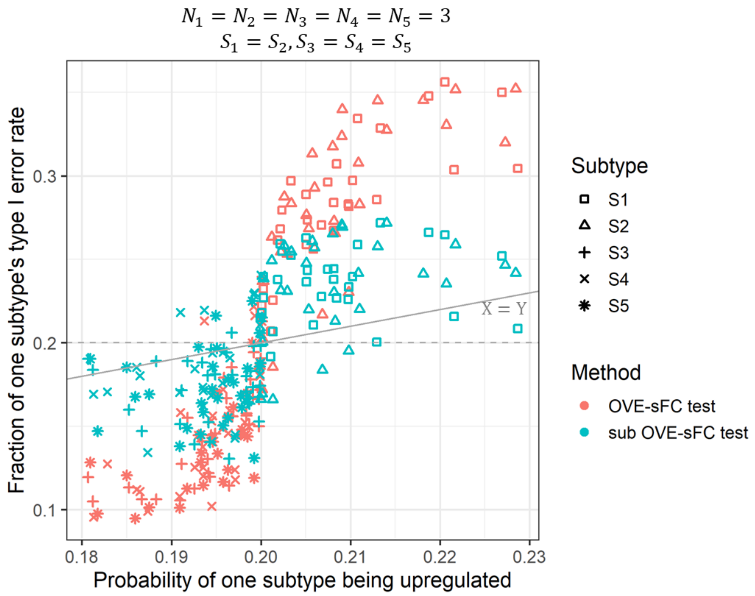


**Figure S4**. Fraction of type I error in each of five subtypes, versus probability of each subtype being upregulated under $H_{\text{non-SDEG}}$ (estimated by Eq. S9). Each point is associated with one of the simulation settings (housekeeping genes: 95%, 80%, 60%, 40%, or 25%; $\sigma_0$: 0.2, 0.5, or 0.8; $\nu_0$: 5, or 40). Sample size is three per subtype. The first two subtypes are drawn from the same one population and the remaining three drawn from another.

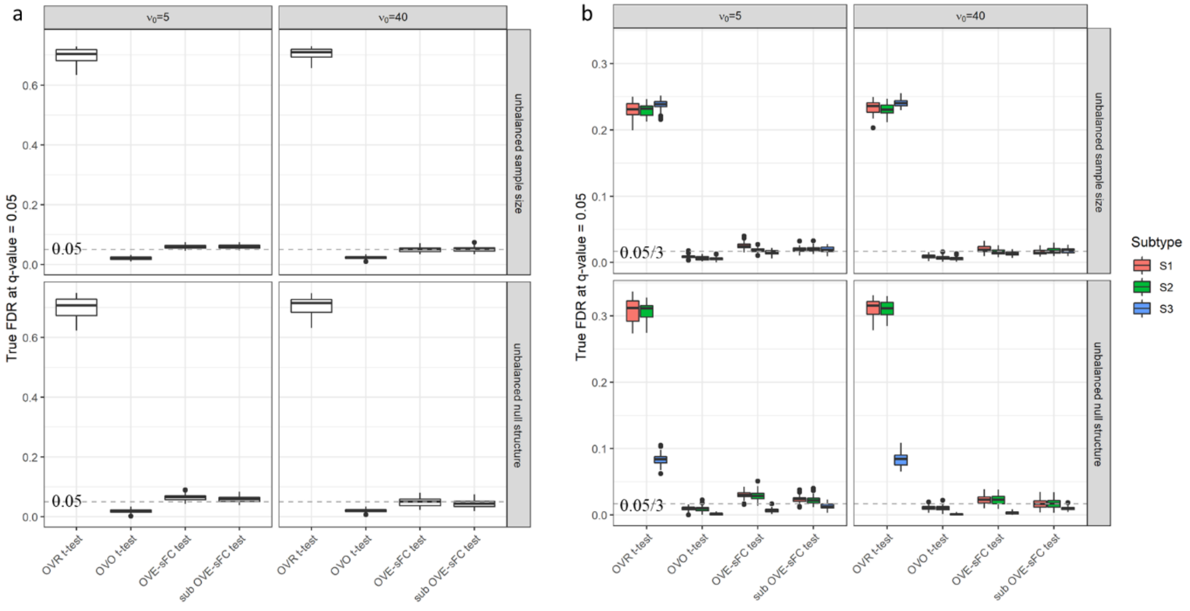


**Figure S5**. FDR control under the multiple simulation settings with three unbalanced subtypes. (a) True FDR at q-value =0.05 across all subtypes (dash line is at 0.05); (b) True FDR at q-value =0.05 in each subtype (dash line is at 0.05/3).

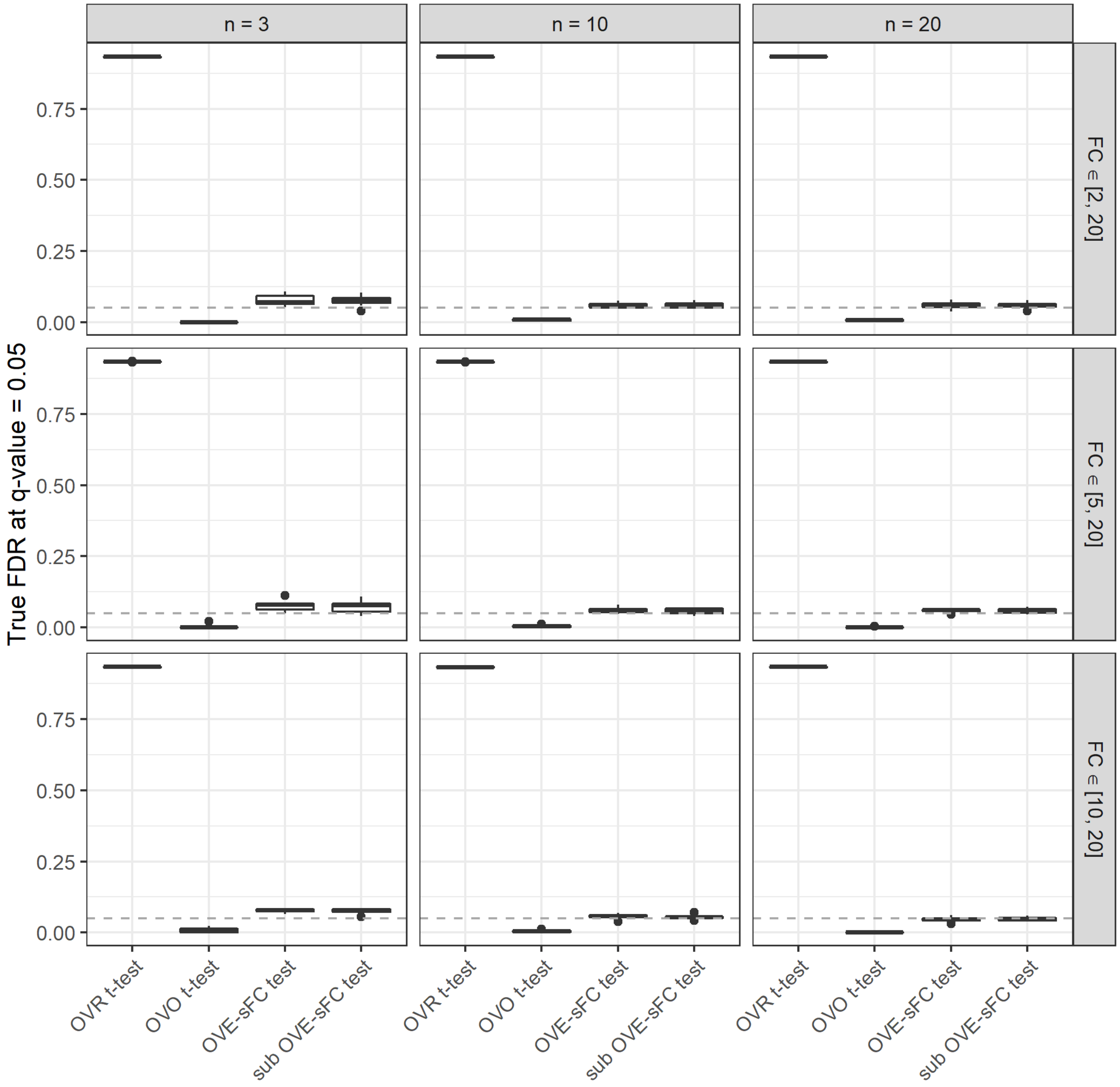


**Figure S6**. FDR control under the multiple simulation settings in RNAseq-derived simulations (dash line is at 0.05). Sample size is 3, 10 or 20 per subtype. Fold change range of SDEGs is [2,20], [5,20], or [10,20]. Non-SDEG distribution is consistent with the base real dataset under the the null hypothesis. Each simulation setting is repeated 20 times

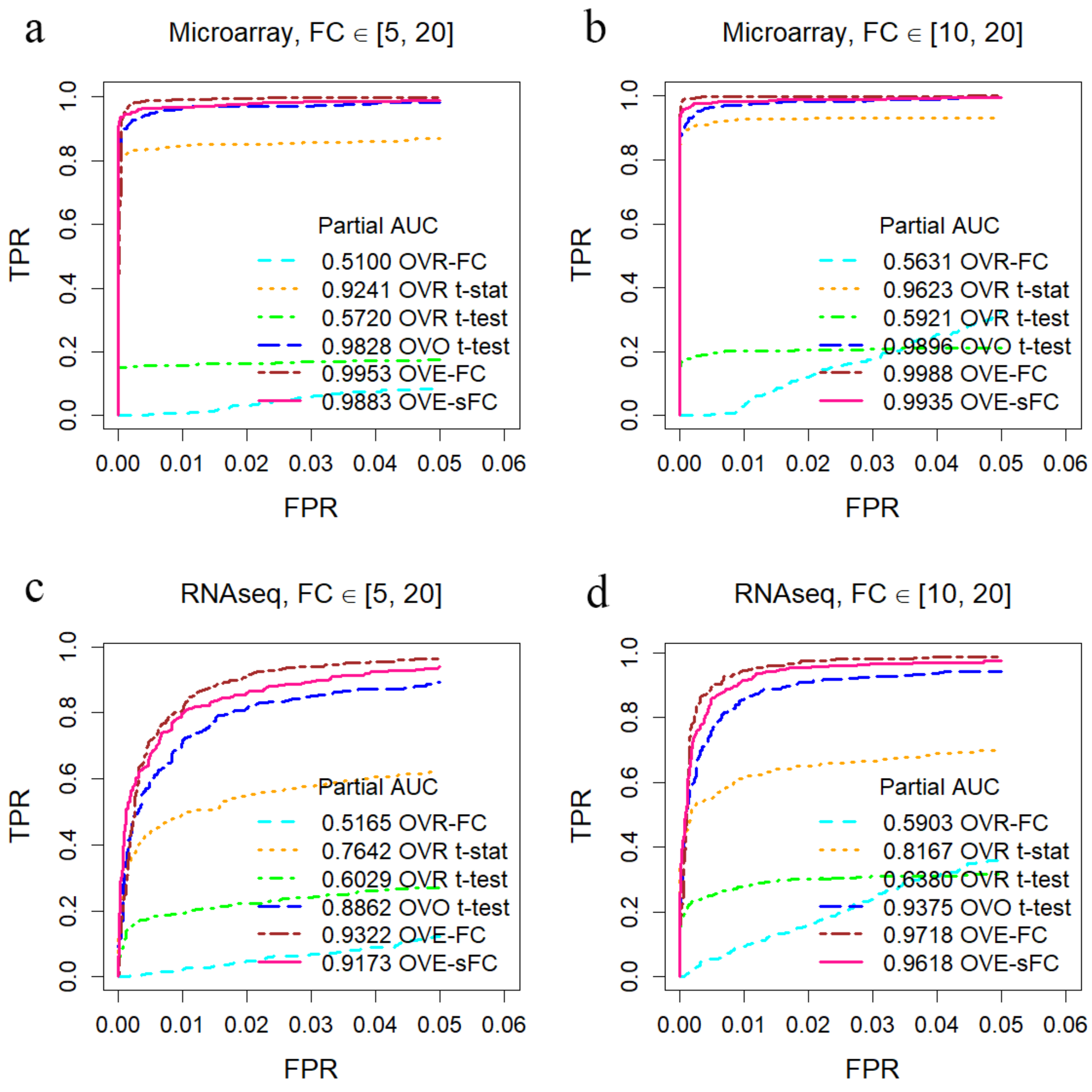


**Figure S7**. Assessment on detection power (partial ROC curves, FPR < 0.05) using real data derived simulations (data distribution is consistent with the base real dataset under the the null hypothesis) involving seven unbalanced subtypes with various parameter settings. Sample size is 3 per subtype. (a)(b) partial ROC curves across different FPR points on microarray-derived data. (c)(d) partial ROC curves across different FPR points on RNAseq-derived data.

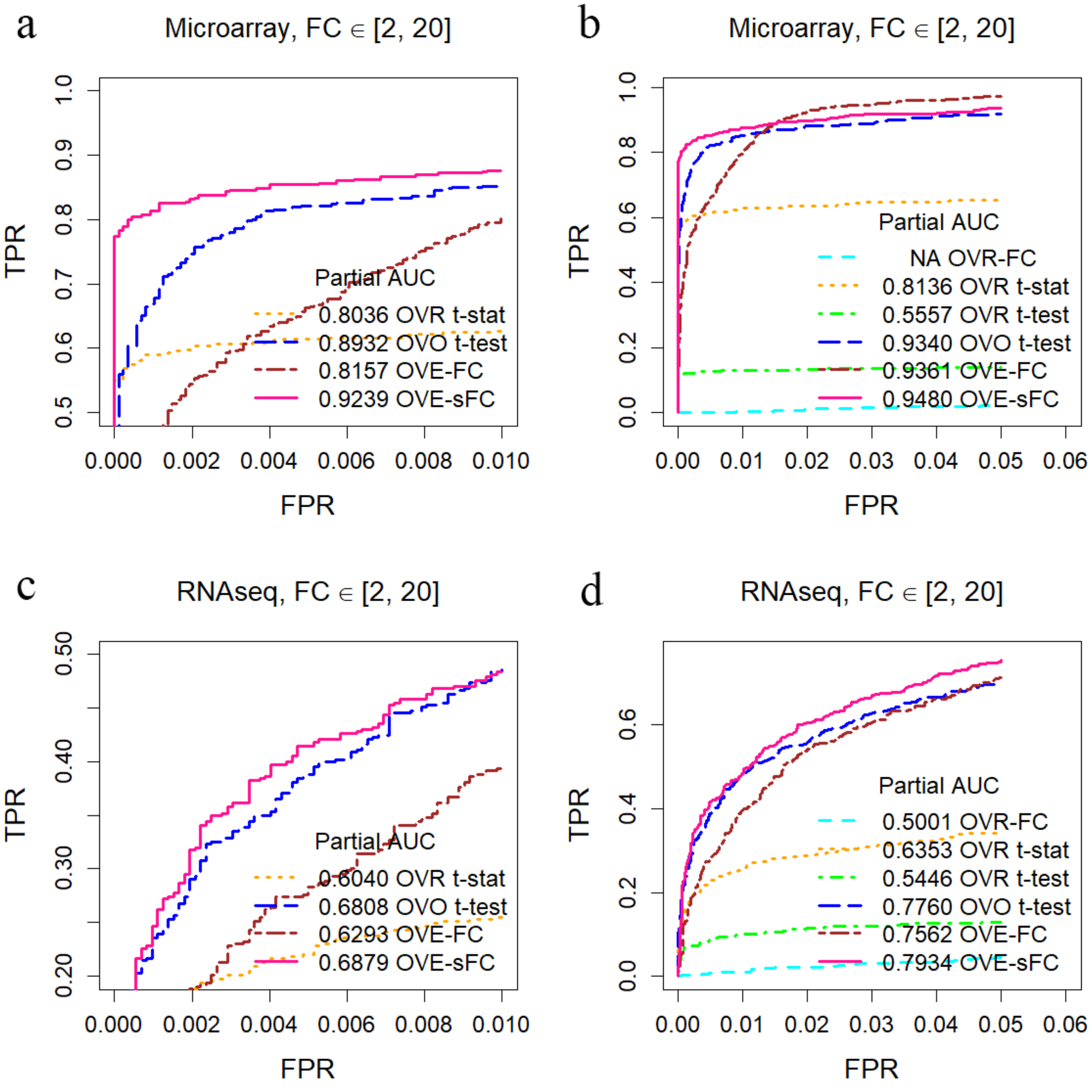


**Figure S8**. Assessment on detection power (partial ROC curves, FPR < 0.01 / 0.05) using real data derived simulations (data distribution is consistent with the base real dataset under the the null hypothesis) involving seven unbalanced subtypes and more nonideal SDEGs with smaller fold change. Sample size is 3 per subtype. (a)(b) partial ROC curves across different FPR points on microarray-derived data. (c)(d) partial ROC curves across different FPR points on RNAseq-derived data.

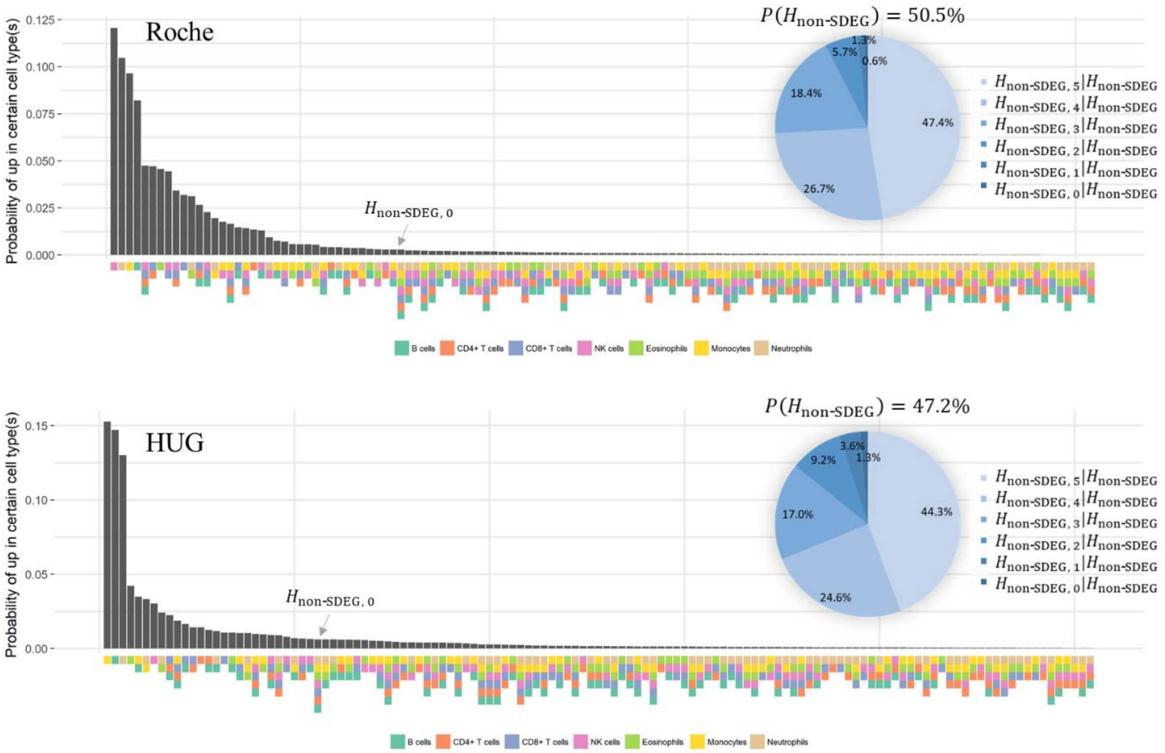


**Figure S9**. Landscape of 127 ($= 2^7 - 1$) gene expression patterns. Estimated probabilities of exclusive expression in any certain cell type(s) are ordered in decreasing value. The pie charts show the conditional probability of each the null hypothesis.

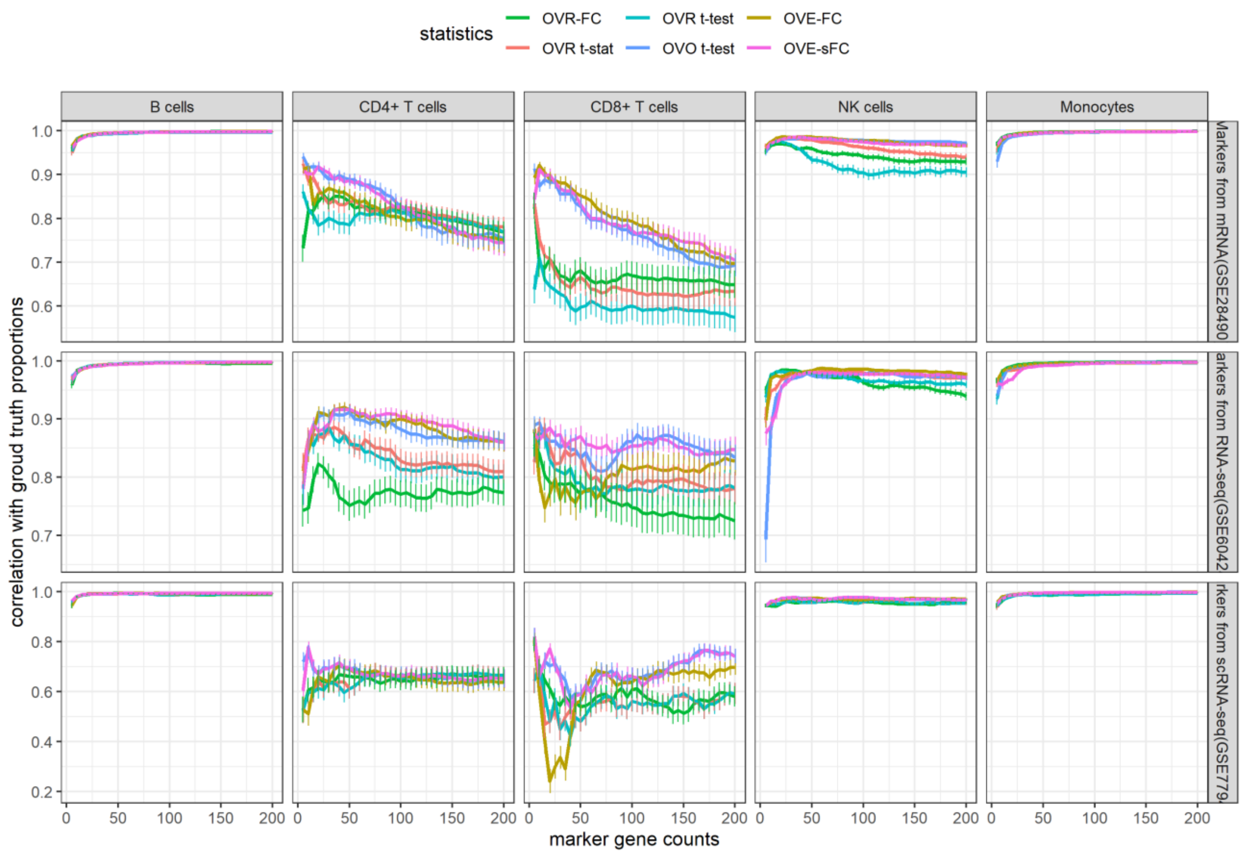


**Figure S10**. Correlation coefficients between CAM score and ground truth proportion for each cell type, with score estimated by a fixed number of markers from independent dataset to quantify subtypes in heterogeneous samples simulated by mixing purified mRNA expression levels in GSE28491. Mean and 95% confidence interval are computed among 20 repeated experiments.

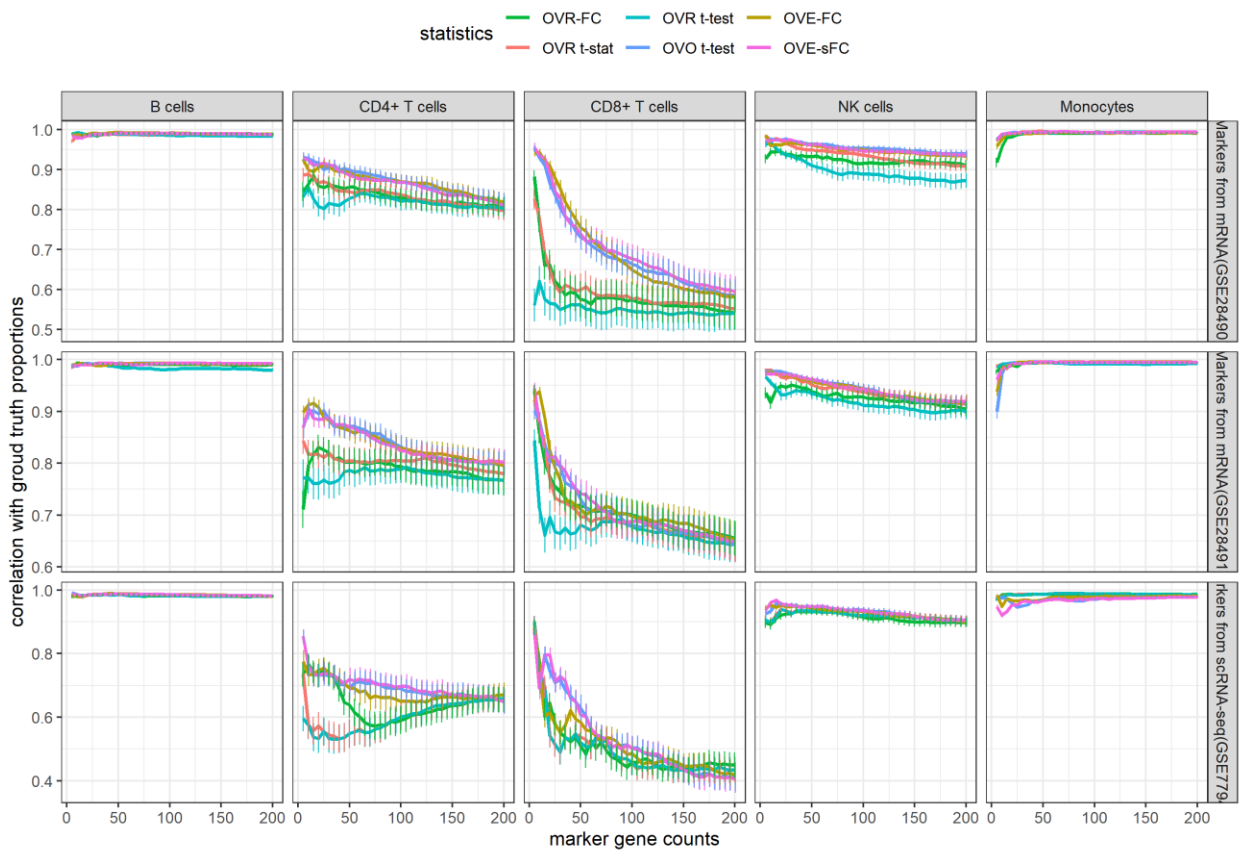


**Figure S11.** Correlation coefficients between CAM score and ground truth proportion for each cell type, with score estimated by a fixed number of markers from independent dataset to quantify subtypes in heterogeneous samples simulated by mixing purified RNAseq counts in GSE60424. Mean and 95% confidence interval are computed among 20 repeated experiments.
